# MIIST305 mitigates gastrointestinal acute radiation syndrome injury and ameliorates radiation-induced gut microbiome dysbiosis

**DOI:** 10.1101/2024.10.22.619652

**Authors:** Debmalya Mitra, Gabriel K. Armijo, Elizabeth H. Ober, Shenda M. Baker, Helen C. Turner, Constantinos G. Broustas

## Abstract

High-dose radiation exposure results in gastrointestinal (GI) acute radiation syndrome identified by the destruction of mucosal layer, intestinal epithelial barrier dysfunction, and aberrant inflammatory responses. In addition, radiation causes gut microbiome dysbiosis characterized by diminished microbial diversity, reduction in the abundance of beneficial commensal bacteria, and the spread of bacterial pathogens that trigger the recruitment of immune cells and the production of pro-inflammatory factors that lead to further GI tissue damage. Currently, there are no FDA-approved countermeasures that can treat radiation-induced GI injury. To meet this critical need, Synedgen *Inc*., has developed a glycopolymer radiomitigator (MIIST305) that is specifically targeted to the GI tract which acts by intercalating into the mucus layer and the glycocalyx of intestinal epithelial cells that could potentially ameliorate the deleterious effects of radiation. Male C57BL/6J adult mice were exposed to 13 Gy total body X-irradiation with 5% bone marrow shielding and MIIST305 was administered on days 1, 3, and 5 post-irradiation. Approximately 85% of the animals survived the irradiation exposure and were apparently healthy until the end of the 30-day study period. In contrast, no control, vehicle-treated animals survived past day 10 at this radiation dose. We show that MIIST305 improved intestinal epithelial barrier function and suppressed systemic inflammatory response mediated by radiation-induced pro-inflammatory cytokines. Taxonomic profiling and community structure of the fecal and colonic mucosa microbiota demonstrated that MIIST305 treatment increased microbial diversity and restored abundance of beneficial commensal bacteria, including *Lactobacillus* and *Bifidobacterium* genera, while suppressing potentially pathogenic bacteria compared with vehicle-treated animals. In summary, MIIST305 is a novel GI-targeted therapeutic that greatly enhances survival in mice exposed to lethal radiation and protects the GI tract from injury by restoring a balanced gut microbiota and effectively reducing proinflammatory responses. Further development of this drug as an FDA-approved medical countermeasure will be of critical importance in the event of a radiation public health emergency.

## Introduction

Accidental or intentional exposure to high-dose radiation often leads to severe hemorrhage, multi-organ failure, and infection, sepsis, and death^1^. The hematopoietic system and the gastrointestinal (GI) tract are particularly vulnerable to radiation-induced damage^2,3^. High-dose radiation exposure can trigger a GI subsyndrome characterized by disruption of the mucosal layer, intestinal epithelial barrier dysfunction, and aberrant inflammatory responses potentially leading to rapid death^3^. While advancements have been made to counteract the immediate effects of hematopoietic acute radiation syndrome^1^, no FDA-approved medical countermeasures (MCMs) currently exist that can treat radiation-induced GI injury^4^.

The lower GI tract harbors a rich community of microorganisms that reside within the luminal and mucosal compartment, which represent distinct niches differing in microbial diversity and composition and play crucial roles in intestinal physiology^5^. A diverse and healthy community of commensal bacteria regulates key epithelial cell functions, contributing to the maintenance of intestinal epithelial barrier integrity and modulating host’s inflammatory state under both homeostatic conditions and in response to injury^6^. In contrast, microbiome dysbiosis promotes aberrant immune responses and susceptibility to inflammatory diseases^7^. The diversity and composition of bacterial flora in the small and large intestine undergoes significant alterations in response to a single, high-dose, irradiation and, conversely, luminal microbiota composition influences the host’s intestinal response to radiation^8–11^ and contributes to increased sensitivity of the gut to inflammation^12^. In rodent studies of high dose radiation exposure (5-18 Gy), specific alterations in the microbiota have been observed which include increased abundance of the phylum Proteobacteria, family Lactobacillaceae, Muribaculaceae and Prevotellacea alongside with decreased abundance of families Lachnospiraceae, Ruminococcaceae, and Clostridiaceae^13–18^

The intestinal epithelial barrier is protected from commensal and pathologic bacteria by three defense mechanisms: the mucus layer^19–20^, the glycocalyx, a 3D matrix rich in carbohydrates and transmembrane mucins, on the apical surface of intestinal epithelial cells^21^ (IECs), and the tight junctions^22^ between intestinal epithelial cells. The mucus layer acts as a first line of defense by preventing luminal bacteria from interacting with the intestinal barrier, thus reducing bacterial exposure of epithelial and immune cells^23^. The primary component of the intestinal mucus is gel-forming, O-linked glycosylated Muc2 polymers, which are predominantly secreted by goblet cells and the gut microbiome has the capacity to alter the production of mucus^24^. These Muc2 polymers are essential for maintaining a stable microbial community in the gut^25^ and support the expression of tight junctional proteins^26^. Goblet cell depletion leads to mucus layer impairment, infection, inflammation and aberrant cytokine response in the gut further leading to improper expression of tight junction proteins that appear to be a component of several gastrointestinal diseases^27^. Even if a pathogenic microorganism manages to penetrate the mucus layer and reach the IEC the surface glycocalyx, will physically inhibit contact between microbe and IECs^28–29^.

With the hypothesis that the glycocalyx is a key therapeutic target for GI-ARS and to meet the critical need for GI-specific MCMs, Synedgen Inc., has developed an orally delivered glycopolymer radiomitigator (MIIST305) that is specifically targeted to the GI tract to potentially alleviate the deleterious effects of radiation. MIIST305 is a polycationic glycopolymer from Synedgen’s multivalent innate immune signaling target platform (MIIST). As a GI-surface targeted therapeutic, MIIST305 associates with the mucus and epithelial glycocalyx^30–31^. These anionic glycopolymers physically impedes a vast majority of microorganisms from accessing the epithelial barrier^28–29^, thus maintaining tolerance towards the commensal gut microbiota. The drug is administered orally and is not absorbed systematically, thus limiting the potential for possible systemic side-effects. Further, extensive non-clinical toxicology and safety pharmacology studies in two animal models have shown no adverse events at maximum deliverable doses of the drug.

In this study we examined the efficacy of MIIST305 to alleviate GI-ARS pathophysiology in male C57BL/6J mice. We assessed animal survival up to 30 days following a single high-dose radiation exposure and showed that MIIST305 significantly reduced mortality. To simulate a realistic radiation exposure scenario, mice were irradiated with 5% bone marrow shielding (BM5), which approximates inhomogeneous radiation exposure likely to occur in a real-world nuclear event, where random shielding would be provided by structures such as buildings^32–33^. Furthermore, we investigated histological injury, intestinal barrier function, and systemic and local cytokine production after radiation exposure. Finally, we analyzed the impact of radiation on the gut luminal and mucosal microbiome diversity and composition in MIIST305 or vehicle-treated mice.

## Materials and Methods

### Ethics statement

The animal studies were reviewed and approved by the Columbia University Irving Medical Center-Institutional Animal Care and Use Committee (CUIMC-IACUC), and all experimental procedure were conducted in accordance with CUIMC-IACUC guidelines and regulations (Protocol no. AABU0652).

### Animals

Eight-week-old male C57BL/6J mice were obtained from Jackson Laboratory (Bar Harbor, ME,) and housed in the animal facilities at CUIMC under specific pathogen-free conditions and five animals were kept per individually ventilated cages. Mice were allowed to acclimate for 10 days prior to experimentation. The animal rooms were maintained at a temperature of 20 ± 3°C and a humidity of 40–60%, with a 12-hour light/dark cycle. The mice were provided with 5053 Irradiated PicoLab^®^ Rodent Diet 20 (Lab Diet ^®^, Arden Hills, MN) and water ad libitum. Wet food pellets and hydrogel packs (Clear H_2_O, West brook, ME) were introduced to the cage floor following irradiation. Animals were randomly assigned into four treatment groups: (1) Vehicle unirradiated (UI), (2) MIIST305 UI, (3) Vehicle irradiated (IR), and (4) MIIST305 IR. Animals were identified by ear punches and cage labels.

### Irradiation

The animals from Vehicle IR and MIIST305 IR groups were irradiated with X-rays using an X- RAD320 irradiator (Precision, Madison, CT), operated at 320 kV,12.5 mA. The table height was set at 52 cm, and the X-rays were administered at doses of 13.0 Gy and 12.5 Gy, with a dose rate of approximately 2.0 Gy/min as detailed in table 1. Dosimetry was performed using a Radcal ion chamber (Monrovia, CA) to monitor the dose rate. Mice were sedated with isoflurane (Covetrus, Portland, ME) and placed in holding fixtures with the left hind leg extended out^34^ and shielded with a 0.6 cm thick lead block covering the femur, tibia, fibula, and paw, resulting in BM5^35^. All irradiations were completed between 10:00 am - 12:00 pm to avoid confounding circadian conditions^36^. The mice from Vehicle UI and MIIST305 UI were sham irradiated.

**Table 1:** Comprehensive overview of irradiation procedure employed in the experimental study.

|  |  |
| --- | --- |
| **Irradiation** |  |
| Instrument | X-RAD320 (320 kV, 12.5 mA) |
| Radiation dose | 13.0 Gy |
| Radiation dose rate | 2.0 ± 0.1 Gy/min |
| Filter | 1 (2 mm Al) |
| Dose rate calibration | RadCal ionization chamber (10X6-6) connected to an Accu-Dose+ digitizer module. |
| SSD | 52 cm |
| PBI | 5 % |
| Mice (male C57BL/6J, 8-10 wks) | Anesthetized by isoflurane |
| **MIIST305 treatment** |  |
| Concentration/mouse | 50 mg/kg delivered in 100 µL water |
| Control mice | 100 µL sterile purified water |
| Delivery route | Oral gavage |
| Dosing | Days 1, 3, 5 post-irradiations |

### Administration of MIIST305

Mice from the MIIST305 (UI/IR) groups were administered MIIST305 (50 mg/kg/day) diluted in sterile endotoxin-free purified water via oral gavage, starting 24 hours post irradiation with two further treatments on days 3 and 5. The vehicle groups received only purified water. The volume of the dosage administered to each mouse was 100 µl.

### Health monitoring

The body weights of the animals were recorded prior to radiation exposure, after which they were returned to their respective cages and monitored daily until 12 days post irradiation (DPI) and once every other day thereafter for signs of body weight loss. Animals that lost 35% of their body weight or displayed signs of morbidity according to the Mouse Interventional Scoring System

(MISS)^37^ were euthanized via CO_2_ inhalation followed by cervical dislocation, in accordance with institutional IACUC guidelines.

### Collection of blood and tissue sample

Blood samples for serum isolation were collected by cardiac puncture along with luminal and mucosal samples at 6 and 12 DPI timepoints for further analysis. A minimum of 5 mice/treatment group were considered for each timepoint.

### Histopathology

The colon was dissected from the small intestine at the ileocecal junction, washed in ice-cold phosphate buffered saline (PBS), and the length was measured in centimeters for each experimental group^38^. To evaluate the severity of colonic damage and inflammation, colon sections were prepared as Swiss rolls and fixed in a 10% neutral-buffered formalin solution for 24 hours, followed by incubation with 70% ethanol. Tissue samples were submitted to the CUIMC Molecular Pathology Core Facility for sectioning and staining with eosin and hematoxylin for microscopic examination^39^ (ECHO revolve, Padgett Street, CA). Tissue sections were assessed for crypt cell proliferation and goblet cell loss.

### Immunohistochemistry (Ki67)

Tissue samples were submitted to the CUIMC-Molecular Pathology Core Facility for Ki67 staining. Briefly, colon tissue samples embedded in paraffin^40^ were dewaxed using xylene and subsequently hydrated. Antigen retrieval was performed, and endogenous peroxidase activity was inhibited with 3% H_2_O_2_. Slides were blocked with 5% BSA blocking buffer at room temperature for 25 min. Subsequently, tissues were incubated with Ki67 polyclonal (Cell Signaling Technology, Inc, Danvers, MA) antibody at 37°C for 2 hours followed by anti-rabbit IgG incubation at 37°C for 1.5 hours. Immunostaining was visualized using DAB substrate solution (Dako, Agilent, Santa Clare, CA), and samples were counterstained with hematoxylin at room temperature for 10–30 seconds^41^. The slides were examined using a light microscope (10X objective lens), and the average number of crypts per field and the number of Ki67-positive cells per crypt was counted across five fields per slide for comparative analysis among the test groups.

### Mucin staining by alcian blue

Dewaxed and rehydrated colonic tissue samples were stained using the Alcian Blue Stain Kit (Vector Laboratories, Newark, CA) according to the manufacturer’s protocol and examined microscopically at (10 x objective)^42^. The intensity of Alcian Blue staining was quantified by assessing the mean gray scale using Image J (NIH, Bethesda, MD). Additionally, the average number of goblet cells was counted across five fields per slide for comparative analysis among the experimental groups^43^.

### Intestinal permeability assay

Test animals were fasted for 4 hours, followed by oral gavage administration of FITC-dextran 4 kDa (Sigma-Aldrich, St. Louis, MO) at a dose of 44 mg/100 g body weight to the experimental mice from the Vehicle IR and MIIST305 IR groups. Mice from the Vehicle UI group received PBS. After 4 hours, mice were euthanized and whole blood was collected by cardiac puncture. The FITC concentration in the serum was quantified fluorometrically using a multimode reader (Synergy H1, Agilent technologies, Santa Clara, CA) at an emission wavelength of 528 nm and an excitation wavelength of 485 nm. All concentrations were measured against a standard curve of serially diluted FITC-dextran^44^.

### Bacterial Translocation assay

To assess bacterial translocation following PBI/BM5 (12.5 Gy), mesenteric lymph nodes (MLN) were aseptically collected from the sacrificed mice of the test groups. Collected MLN tissues (1 mg) were homogenized in 1ml PBS. Homogenates were plated on MacConkey agar plates (BD DIFCO, Franklin Lakes, NJ) and incubated at 37°C for 24–48 hrs. Colony forming units (CFU) were counted, and densities calculated as follows: CFU/mL = (number of colonies × sample dilution factor × serial dilution factor)/volume of culture plate (mL)^45–46^.

### Cytokine analysis

Cytokine quantification in blood serum and colonic tissue samples was performed using the U-PLEX assay from Meso Scale Discovery multiplex arrays (MSD, Rockville, MD). Serum samples were diluted two-fold according to the manufacturer’s instructions, and cytokine concentration levels were reported in pg/mL of serum. Cytokine levels in colonic tissue were expressed as pg/100 mg of tissue^47^. Briefly, flash-frozen colonic samples were homogenized in ice-cold cell lysis buffer (50 mM Tris, pH 7.4, 250 mM NaCl, 5 mM EDTA, 50 mM NaF, 1 mM Na_3_VO_4_, 1% NP- 40, 0.02% NaN_3_) containing 2X protease inhibitor, using three 40-second bursts using Mini Bead beater 16 (Biospec Products, Bartlesville, OK). The homogenate was then centrifuged at 14,000 rpm for 10 minutes at 4°C. The supernatant was collected and used for the assay after determining protein concentrations using the BCA assay method (Pierce, Thermo Fisher Waltham, WA).

### Microbiome analysis

Stool and colonic tissue samples from mice were collected, flushed with PBS, frozen in liquid nitrogen, and stored at -80°C. Samples were sent to Zymo Research (Irvine, CA) on dry ice for DNA extraction, library preparation, and sequencing. DNA extraction was performed using the ZymoBIOMICS^®^ DNA Miniprep kit, and targeted sequencing was prepared with the Quick-16S™ Plus NGS Library Prep Kit (V3-V4 Primer Set). ZymoBIOMICS^®^ Microbial Community DNA Standard was used as a positive control. Real-time PCR was employed to control cycles and minimize PCR chimera formation. Final PCR products were quantified by qPCR, pooled by equal molarity, and cleaned with Select-a-Size DNA Clean & Concentrator™. The library was quantified with TapeStation^®^ (Agilent Technologies, Santa Clara, CA) and Invitrogen Qubit 1X dsDNA High-Sensitivity Assay Kits^®^ (Thermo Fisher Scientific, Waltham, WA) and sequenced on Illumina^®^ NextSeq 2000™ (600 cycles) with 30% PhiX spike-in. DADA2 pipeline was used to infer unique amplicon sequences and remove chimeras^48^. Taxonomy assignment was performed using UCLUST from QIIME v1.9.1, with biom files analyzed in Microbiome Analyst 2.0^49^ using default parameters: minimum count of 4, 20% prevalence, inter-quantile range for low variance filter, total sum scaling for normalization, and no rarefaction or transformation. α-diversity was assessed using Chao1 and Shannon indexes, and β-diversity via Bray-Curtis dissimilarity with PCoA visualization. Differential abundance was calculated using DEseq2.

### Statistical Analysis

Statistical analyses were performed using GraphPad Prism 10. Kaplan-Meier curves, accompanied by log-rank tests, were employed for survival analysis^32^. For histological analysis and cytokine analysis, comparisons between Vehicle and MIIST305 were made using two-tailed unpaired t-test^50^. Data are presented as mean ± SEM and results were considered statistically significant at P < 5.0E-02. For microbiome analysis, α-diversity was assessed using the (Mann-Whitney U test for two groups, and the Kruskal-Wallis multiple comparison test for more than two groups. β-diversity was evaluated with PERMANOVA (Permutational multivariate analysis of variance)^51^. Differential abundance analysis of specific genera was determined by DESeq2 and a p < 0.05 (nonparametric test Wilcoxon rank-sum test) was considered significant.

## Results

### MIIST305 increases survival of experimental mice following a lethal dose of acute radiation exposure

To evaluate the mitigating efficacy of MIIST305, Vehicle and MIIST305-treated C57BL/6 male adult mice (n = 20/group) were exposed to 13 Gy of PBI/BM5, and survival was determined up to 30 days post-irradiation. Unirradiated mice (n = 10/group) treated with MIIST305, or vehicle were used as controls. Exposing vehicle-treated mice to 13 Gy X-rays was lethal to 100% of vehicle-treated mice by day 10 following irradiation, whereas approximately 85% (p < 1.0E-03) of MIIST305-treated animals survived 30 days post-irradiation (Figure 1a). In both vehicle- and MIIST305-treated cohorts, animal death occurred between days 7 and 10, consistent with the GI-ARS time course. All unirradiated mice survived until the end of the experiment on day 30. Both vehicle and MIIST305-treated animals lost weight in response to 13 Gy X-rays at comparable rates for the first 6 days (Figure 1b). After that, the weight loss of the MIIST305 irradiated mice was significantly less (p < 5.0E-02) compared with vehicle-treated animals, reaching a nadir of approximately 30% on day 8 and recovery starting from day 9 onwards. By day 14, the MIIST305 mice had regained on average 86% of their pre-irradiation weight that was maintained until the end of the 30-day study period. In contrast, the vehicle-treated irradiated mice continued to lose weight, and by day 10, all surviving animals had to be euthanized either due to excessive weight loss (approximately 35%) or signs of morbidity.

**Figure 1.**
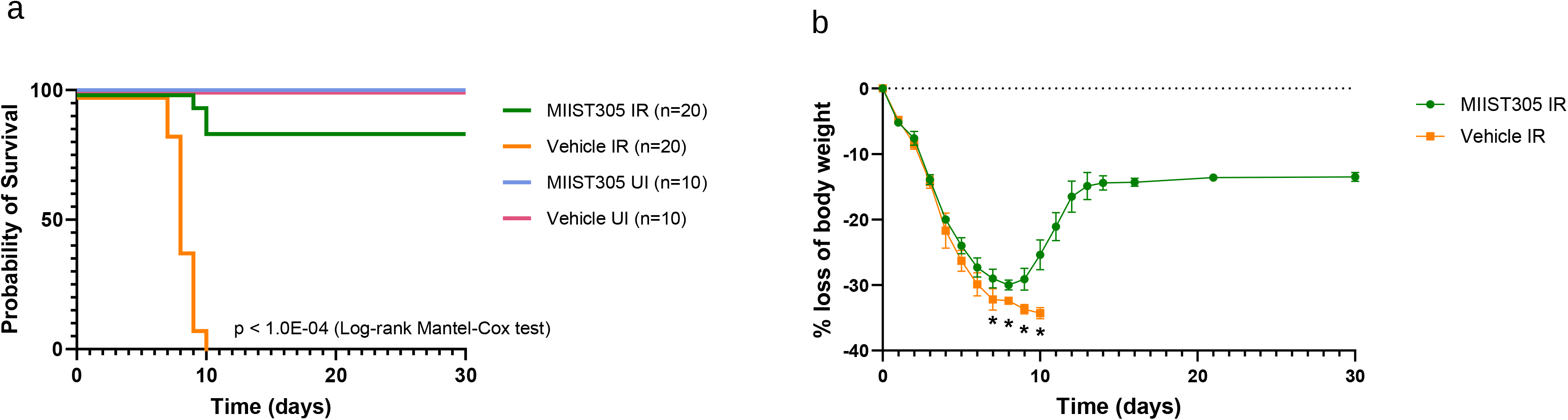
Administration of MIIST305 increases survival and prevents body weight loss in C57BL/6 mice exposed to 13 Gy PBI/BM5. (a) The Kaplan-Meier curve illustrates the survival of the treatment groups up to 30 DPI. Treatment groups were Vehicle UI (n = 10), MIIST305 UI (n = 10), Vehicle IR (n = 20), and MIIST305 IR (n = 20). Statistical analysis was performed by Log- rank (Mantel-Cox) test (b) Comparison of body weight loss (%) following exposure to X-rays at 13.0 Gy, PBI/BM5 measured till 30 DPI (n = 20). Data are presented as mean ± SEM, two-tailed unpaired t-tests were conducted between treatment groups. *p < 5.0E-02 is considered statistically significant with respect to the Vehicle IR.

### Administration of MIIST305 improves radiation-induced enteropathy

To analyze temporal colonic structural and functional changes, systemic and local inflammation, as well as gut microbiome changes following high dose irradiation, we reduced the radiation dose to ensure survival of more of the vehicle-treated animals at later time points after day 10. At a dose of 12.5 Gy PBI/BM5, we observed that 30% of vehicle-treated animals survived past day 21 (Figure S1) and therefore, this radiation dose was used to conduct our subsequent studies.

Radiation exposure resulted in significant colon length shortening in both Vehicle- and MIIST305-treated animals at 6 DPI, which is a sign of severe tissue injury. Figure 2a depicts the mean colon length of Vehicle UI mice (n = 5) was 7.8 ± 0.47 cm, which was significantly reduced by 24.87 ± 1.46 % (p = 2.0E-04) in the Vehicle IR mice (n = 5) at 6 DPI. The MIIST305 IR mice (n = 5) also exhibited a significant reduction in colon length at 6 DPI by 20.25% ± 20.83 % (P = 2.0E- 02) compared to the MIIST305 UI. The colon length between the irradiated Vehicle- and MIIST305-treated mice (n = 15) was similar at 6 DPI with a mean length of 6.14 ± 0.86 cm and 6.53 ± 0.80 cm, respectively. However, by 12 DPI, the mean colon length of the MIIST305 IR mice (n = 10) were found to be 7.37 ± 0.66 cm which was significantly longer by 19.67 ± 3.77 % (P = 1.0E-04) compared to the Vehicle IR mice (n = 15) (Figure 2b), indicating the efficacy of MIIST305 in mitigating irradiation-induced damage to the colon.

**Figure 2.**
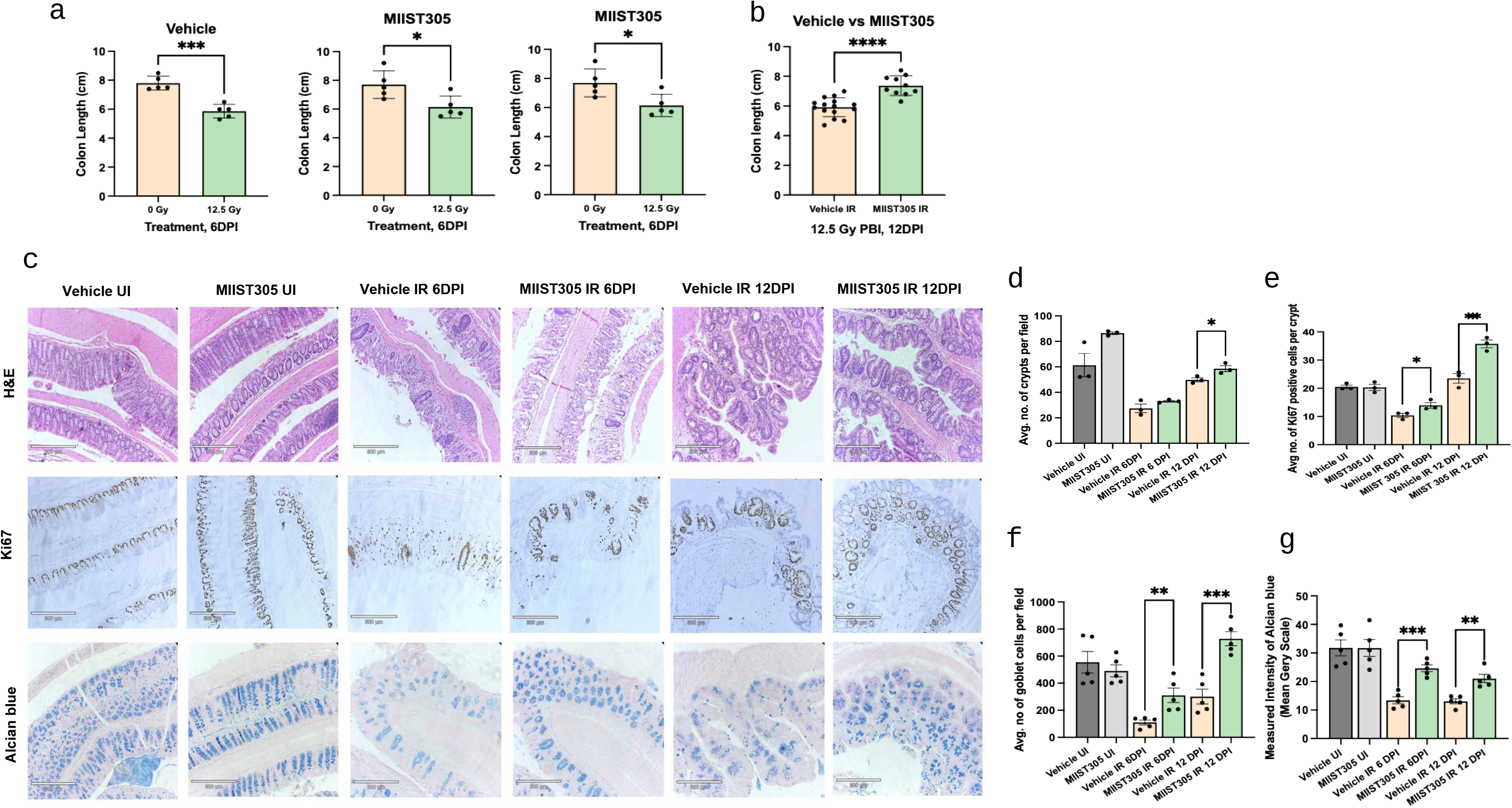
MIIST305 prevents X-ray-induced colon structural and functional damage in C57BL/6 mice at 12 DPI. (a) Colon lengths from treatment groups were measured at different time points: Vehicle UI (n = 5) vs. Vehicle irradiated IR 6 DPI (n = 5); MIIST305 UI (n = 5) vs. MIIST305 IR 6 DPI (n = 5); Vehicle IR 6 DPI (n = 5) vs. MIIST305 IR 6 DPI (n = 15) (b). Vehicle IR 12 DPI (n=15) vs. MIIST305 IR 12 DPI (n=10). (c) histological analysis of colon cross-sections using H&E staining, Ki67 immunohistochemistry, and Alcian blue staining, (10x objective, Scale bar: 50 µm), (d) average number of crypts per field (n = 3), (e) average number of Ki67-positive cells per crypt (n =3 ), (f) Average number of goblet cells per field (n = 5), (g) mean grayscale intensity of Alcian blue staining (n = 5). Comparisons between treatment groups were performed using two-tailed unpaired t-tests, and the data are presented as mean ± SEM. For colon length, Statistical significance is indicated as *p < 5.0E-02, with respect to MIIST305 0 Gy, ***p < 5.0E-04, with respect to Vehicle 0 Gy, ****p < 5.0E-05, with respect to Vehicle IR. For Histological analysis Statistical significance is indicated as *p < 5.0E-02, **p < 5.0E-03 and ***p < 5.0E-04, with respect to Vehicle IR.

Histopathological examination demonstrated loss of colonic crypts on 6 DPI with a partial recovery on 12 DPI, that was higher in the MIIST305-treated mice compared to the Vehicle only mice (Figure 2c and 2d).The average number of crypts in the tissue sections of the Vehicle UI and MIIST305 UI group were 61.33 ± 15.64 and 86.46 ± 1.85, respectively which was reduced by 55.22 ± 62.90 % and 61.70 ± 37.29 % in Vehicle IR and MIIST305 IR at 6 DPI with respect to Vehicle UI and MIIST305 UI. MIIST305 treatment did not have any significant impact on the number of crypts at 6 DPI but at 12 DPI, an increase in the average number of crypts was observed in MIIST305-treated mice by 14.91 ± 31.57 % (p = 3.5E-02) with respect to the Vehicle only mice. The average number of proliferative cells / crypts as determined by Ki67 immunostaining was significantly higher in the MIIST305-treated group by 25.25 ± 29.05 % (p = 4.0E-02) at 6 DPI and by 34.28 ± 17.18 % (p = 3.0E-02) at 12 DPI compared to the Vehicle-only treated group as represented in Figure 2e.

Loss of goblet cells and dysregulation of mucus production and secretion has been associated with intestinal epithelial barrier dysfunction, microbiome dysbiosis, increased inflammation and infectious diseases. Staining with the goblet cell marker Alcian Blue (stains acidic mucins and mucopolysaccharides) demonstrated that mice treated with MIIST305 had a significantly higher number of goblet cells compared with vehicle-treated mice on 6 DPI and 12 DPI (Figure 2c and 2f). The average number of goblet cells per field in the Vehicle UI and MIIST305 UI groups were 554.64 ± 178.75 and 491.08 ± 98.81 respectively, (Figures 2c and 2f). These numbers were significantly reduced by 80.11 ± 78.32 % (p = 6.0E-04) and 36.78 ± 16.73 % (p = 3.0E-02) following 12.5 Gy PBI/BM5 exposure in the Vehicle IR and MIIST305 IR groups respectively at 6 DPI (Figure 2f). However, at 6 DPI and 12 DPI, the average number of goblet cells was significantly higher in the MIIST305-treated group by 64.48 ± 3.39 % (p = 7.1E-03) and 58.65 ± 5.94 % (p = 5.0E-04), respectively compared to the Vehicle IR.

Likewise, mucin production per goblet cells, quantitated by measuring the intensity of the Alcian Blue signal per cell, was also significantly higher in MIIST305-treated animals (Figure 2c and 2g). The mean gray scale of Alcian blue staining was highest in the Vehicle UI (31.76 ± 6.26) and MIIST305 UI (31.73 ± 6.66) groups, as represented in Figure 2g. At 6 DPI and 12 DPI, the mean gray scale of MIIST305 IR was significantly higher by 45.53 ± 0.36 % (p = 2.0E-04) and 37.85 ± 39.81 % (p =1.5E-03) compared to the Vehicle-only treated group.

### MIIST305 reduces intestinal epithelial barrier permeability and bacterial translocation in mice exposed to irradiation

To investigate whether MIIST305 treatment administered at 24 h post – radiation with additional doses on days 3 and 5 improves intestinal epithelial barrier function, we performed the FITC-dextran barrier permeability assay and assessed bacterial translocation from the intestinal lumen to the MLNs on 6 DPI involving animals (n = 3 / group) from the Vehicle UI, Vehicle IR and MIIST305 IR experimental groups. As expected, radiation exposure resulted in a large increase in barrier permeability in vehicle-treated mice compared with unirradiated mice. However, treatment with MIIST305 decreased barrier permeability defects, observed by the lower concentration of FITC-dextran (55.77 ± 42.66 %; P = 4.0E-02) in the serum on 6 DPI compared with the Vehicle IR group (Figure 3a).

**Figure 3.**
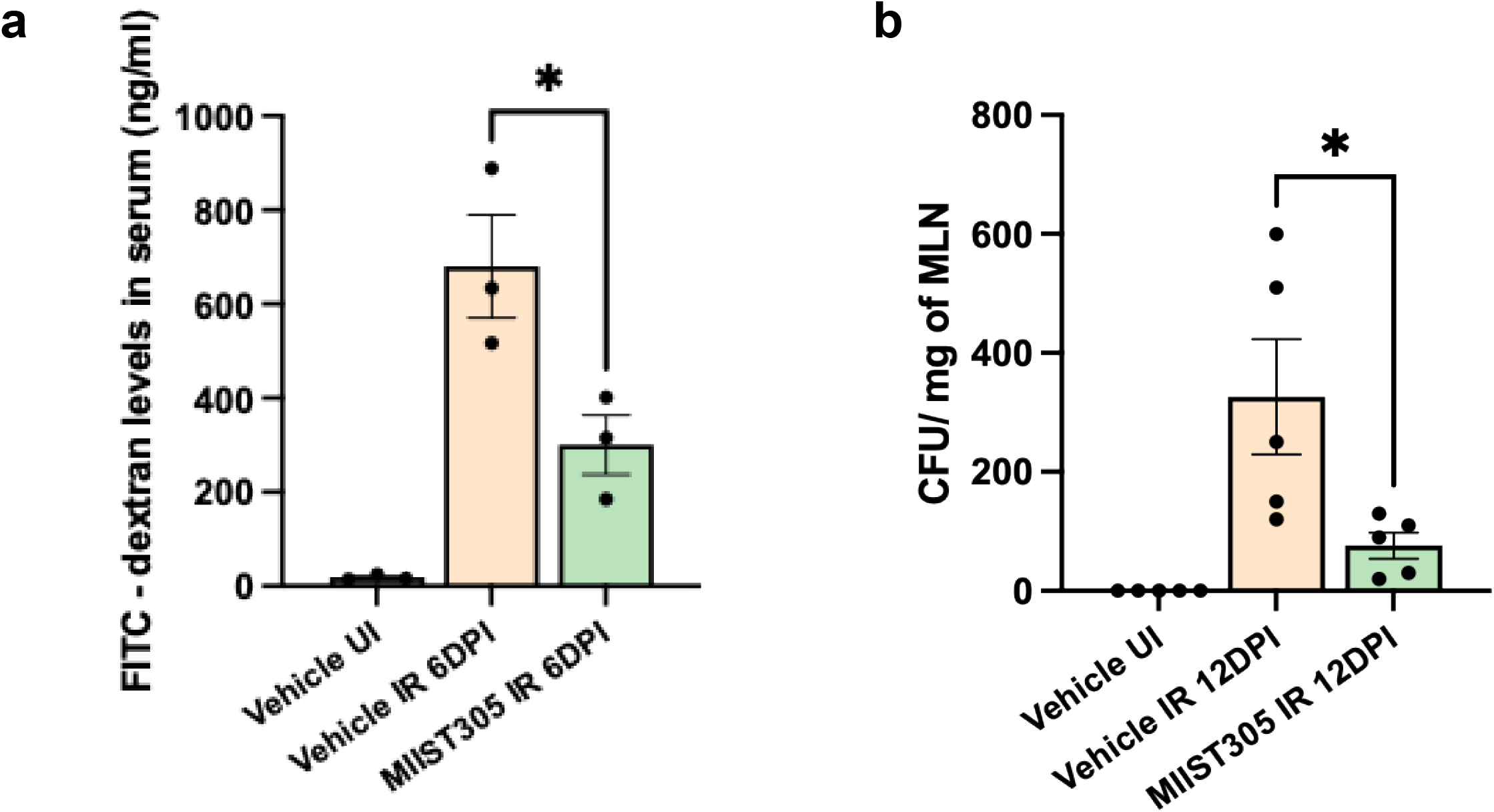
MIIST305 ameliorates gut barrier function. (a) Fluorometric quantification of FITC- Dextran levels (ng/ml) obtained from serum samples of treatment groups (n = 3), (b) Count of CFU on MacConkey agar to quantify gut bacterial translocation in MLN (n = 5). Data are represented as mean ± SEM, two-tailed unpaired t-tests were conducted between treatment groups, *p < 5.0E-02 is considered statistically significant with respect to Vehicle IR (6 DPI/12 DPI).

Bacterial translocation to MLNs indicates a defective gut barrier function. MLNs were collected from Vehicle UI, Vehicle IR and MIIST305 IR groups (n = 5 animals / group) at 12 DPI and obtained result shows that radiation exposure resulted in significant increase in bacterial translocation to MLNs compared with unirradiated mice (Figure 3b). However, treating irradiated mice with MIIST305 significantly inhibited bacterial translocation. Notably, bacterial translocation was significantly reduced in MIIST305-treated mice by 76.68 ± 77.58 % (p =3.0E-02) compared to the Vehicle IR group on 12 DPI.

### Oral administration of MIIST305 modulates serum and colonic cytokine levels following radiation exposure

It is well-documented that radiation exposure elicits a cytokine storm both at the systemic and damaged organ-specific level. Here we measured the levels of a panel of 29 cytokines in both serum and colonic tissue samples (n = 3-8/group) at 6 DPI and 12 DPI following 12.5 Gy PBI/BM5 X-ray exposure. Radiation increased levels of several pro-inflammatory cytokines in the serum including TNF-α, KC/Gro, IP-10, MIP-1β, MIP-2, MIP-3α, MCP-1, IL-17A/F, IL-6, as well as the anti-inflammatory cytokine IL-10 (Figure 4). Specifically, TNF-α and KC/Gro levels were reduced by 25.47 ± 50.66 % (p = 2.0E-02) and 43.44 ± 79.24 % (p = 5.0E-02) in the MIIST305 IR mice when compared to the Vehicle only group at 6 DPI, whereas the expression levels of IP-10, MIP- 1β, MIP-2, MIP-3α, MCP-1 and IL-17A/F were significantly lower in the MIIST305 IR group compared to the Vehicle IR group by 41.29 ± 17.74% (p = 5.0E-02), 54.26 ± 58.15 % (p = 4.0E- 02), 74.68 ± 61.37 % (p = 9.0E-03), 91.21 ± 77.27 % (p = 2.0E-03), 53.49 ± 16.60 % (p = 4.0E-03) and 45.22% ± 69.41% (p = 7.0E-03) respectively, at 12 DPI. No significant change was observed in serum levels of IL-6 and IL-33 in MIIST305 IR at 6 DPI and 12 DPI respectively. In contrast, the concentration of anti-inflammatory cytokine IL-10, was significantly increased in the MIIST305 IR mice by 32.25 ± 57.64 % (p = 2.0E-02) compared to the Vehicle only mice at 12 DPI (Figure 4k).

**Figure 4.**
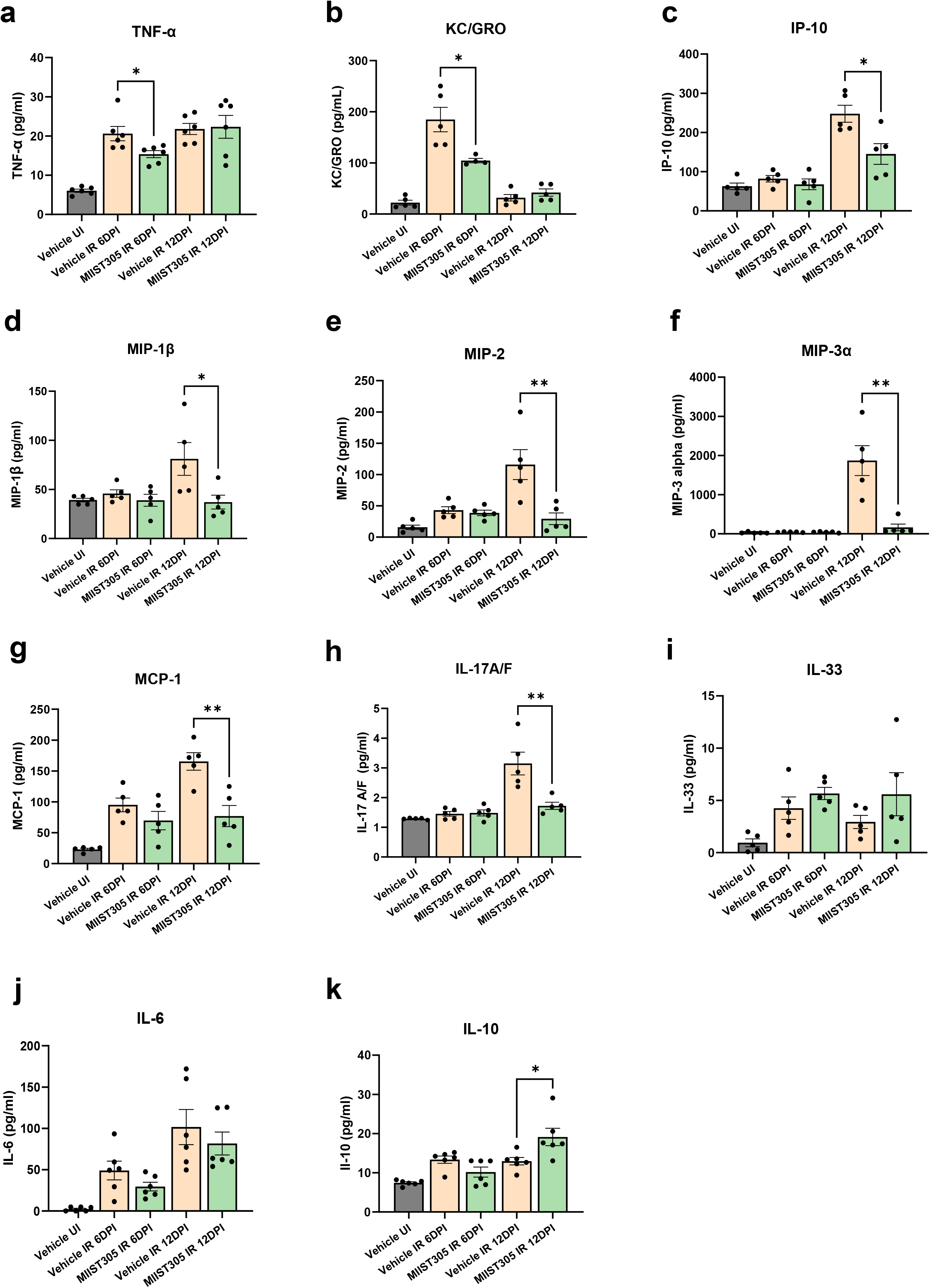
MIIST305 mitigates X-ray-induced inflammation by reducing pro-inflammatory cytokine levels in serum and increasing the anti-inflammatory cytokine IL-10. Quantification of serum cytokines levels (pg/ml) using MSD assay (a) TNF-α, (b) KC/GRO, (c) IP-10, (d) MIP-1β, (e) MIP- 2, (f) MIP-3α, (g) MCP-1, (h) IL-17A/F, (i) IL-33, (j) IL-6 and (k) IL-10. Data are represented as mean ± SEM, two-tailed unpaired t-tests were conducted between treatment groups, *p < 5.0E- 02 is considered statistically significant with respect to Vehicle IR (6DPI/12DPI), **p < 5.0E-03 is considered statistically significant with respect to Vehicle IR (6DPI/12DPI).

In colonic tissue lysates, IL-6 expression was significantly reduced by 48.51 ± 92.44 % (p = 7.0E-03) in MIIST305-treated mice compared to the Vehicle-treated at 6 DPI, whereas IL-33 levels in the MIIST305 IR group increased by 33.54 ± 10.01% (p = 1.8E-02, n = 8) compared to the Vehicle IR group at 12 DPI (Figure 5). Although levels of TNF-α, IP-10, MCP-1, MIP-1β, MIP- 2, and KC/Gro increased in the tissue lysates in response to radiation on 6 and 12 DPI, their levels were similar in both vehicle and MIIST305-treated animals. The expression of IL-17A and IL-22 increased in response to irradiation in serum and tissue lysates but there was no notable difference upon MIIST305 treatment. Further, IL-1β, IL-2, IFN-γ, IL-27p28/IL-30, IL-23 and MIP- 1α, did not respond to irradiation in either serum (Figure S2) or the tissue lysates samples (Figure S3).

**Figure 5.**
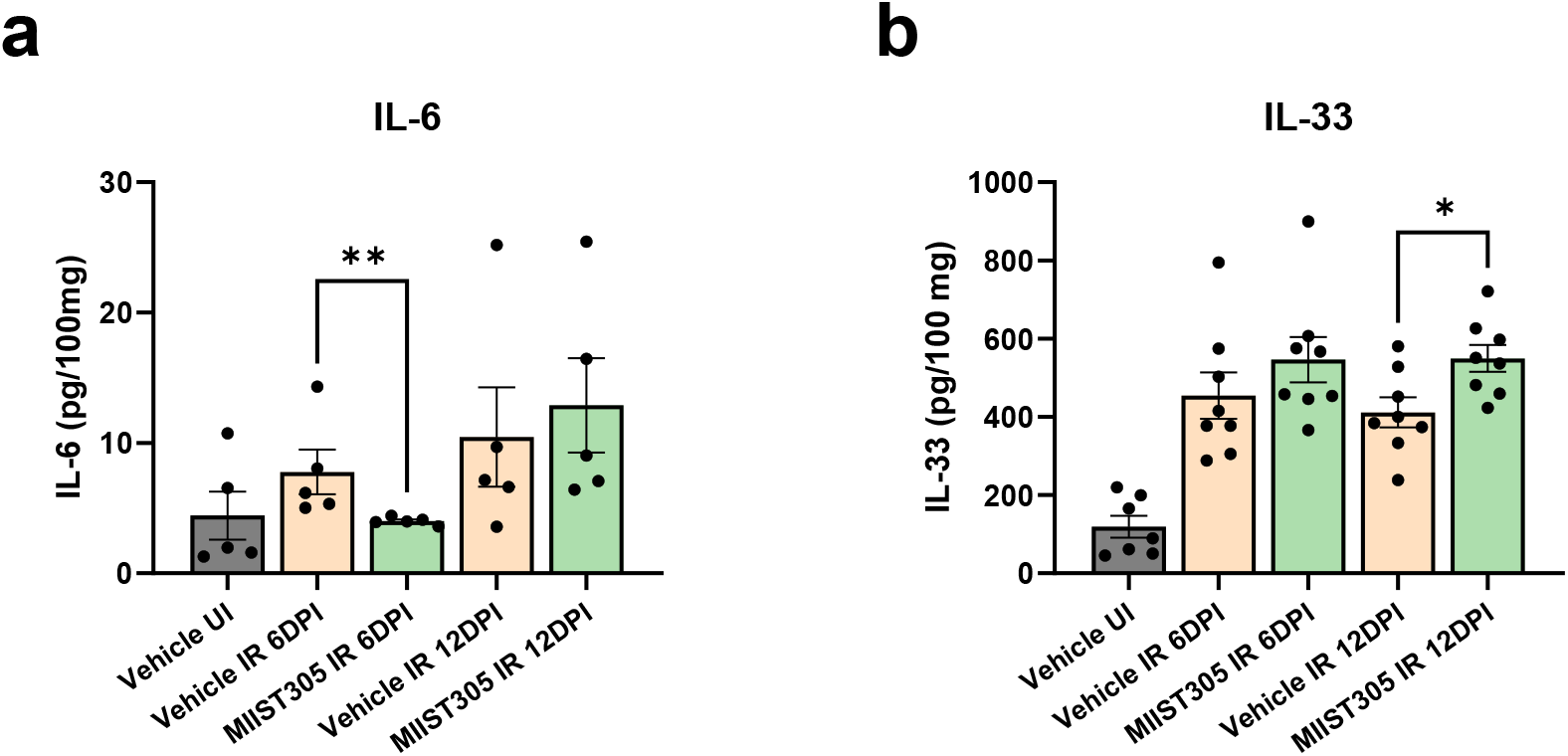
MIIST305 reduces X-ray-induced inflammation by modulating cytokine levels at colonic tissue, specifically decreasing IL-6 and increasing IL-33. Quantification of cytokine levels (pg/100 mg) in colonic tissue was performed using MSD assay for (a) IL-6 and (b) IL-33. Data are presented as mean ± SEM. Two-tailed unpaired t-tests were conducted between treatment groups, with *p < 5.0E-02 considered statistically significant compared to Vehicle IR 12 DPI and **p < 5.0E-03 considered statistically significant compared to Vehicle IR 6 DPI.

### MIIST305 restores a more normal microbiome distribution in GI injury

Gut microbiota reinforces the intestinal epithelial barrier function and reduces inflammation^52^. However, radiation exposure triggers intestinal microbiome dysbiosis affecting the diversity and composition of microbiota that lead to intestinal epithelial cell barrier damage and local and systemic inflammation^53^. To investigate the effect of 12.5 Gy X-ray PBI/BM5 on the gut microbiome of C57BL/6 mice, 16S rRNA amplicon sequencing was conducted on mucosal and luminal samples from the Vehicle- and MIIST305-treated IR mice, analyzed at 6 and 12 DPI. Comparative analysis was performed against unirradiated controls (Vehicle UI and MIIST305 UI). Radiation exposure resulted in a significant reduction in α -diversity (p < 5.0E-02) at the genus rank as judged by the Chao1 index, which accounts for taxa richness within a group, in both the luminal and the mucosal compartment on day 6 (Figure. 6a). Although not statistically significant, mucosal microbiota showed greater α-diversity compared with luminal microbiota. MIIST305 treatment did not have an influence on α-diversity. On the other hand, Shannon index, which takes into consideration the richness and evenness of taxa in the same group (Figure 6b) depicts values that were similar in the unirradiated/irradiated samples and vehicle- and MIIST305-treated samples. Again, mucosal samples displayed higher, but not statistically significant, values compared with luminal samples in both unirradiated and irradiated cohorts. Principal coordinates analysis (PCoA) visualization of β-diversity (Figure 6c – 6h) at the genus level using the Bray-Curtis dissimilarity metric revealed a clear distinction in microbial diversity between Vehicle UI and Vehicle IR 6 DPI in luminal (p = 5.0E-03) and mucosal (p = 1.3E-02) samples. Similar results were also observed with luminal (p = 1.2E-02) and mucosal (p = 9.0E-03) samples of MIIST305 UI and MIIST305 IR 6 DPI. However, at 6 DPI the microbiome profiles of Vehicle IR and MIIST305 IR were closely related in the PCoA plot for both luminal and mucosal samples highlighting their similar composition.

**Figure 6.**
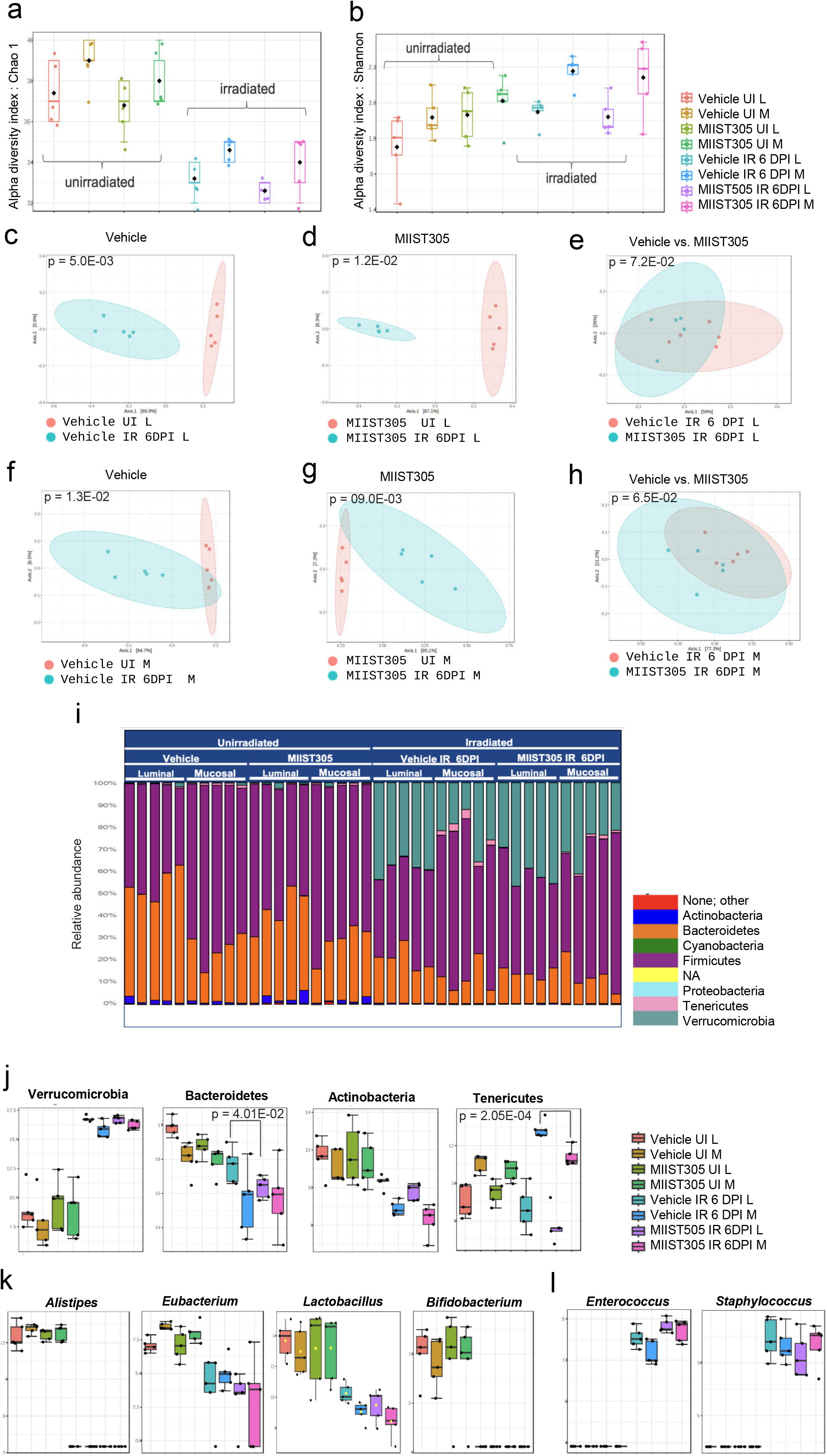
X-ray induces gut dysbiosis and alters gut microbial composition in C57BL/6 (n=5) mice at 6 DPI. Representation of (a) Chao index (b) Shannon index. PCoA analysis of (c) Vehicle UI L vs Vehicle IR 6DPI L, (d) MIIST305 UI L vs MIIST IR 6DPI L, (e) Vehicle IR 6 DPI L vs MIIST305 IR 6DPI L (f) Vehicle UI M vs Vehicle IR 6DPI M, (g) MIIST305 UI M vs MIISt305 IR 6DPI M (h) Vehicle IR 6DPI M vs MIIST305 6DPI M, (i) the relative abundance at phylum level, (j) differential abundance of bacteria at phylum level (k) differential abundance of bacteria at genus levels. L and M denotes luminal and mucosal, respectively. For α-diversity (Chao1 and Shannon index) and β-diversity (PCoA plots) analysis, comparison between test groups was conducted using the Mann-Whitney U test and the PERMANOVA test respectively. For relative abundance at phylum level and differential abundance at genus levels, comparisons between treatment groups were performed using two-tailed unpaired t tests, the data are presented as mean ± SEM and p < 5.0E- 02 is considered statistically significant.

Taxonomic profiling at the phylum level (Figure 6i and 6j) showed that unirradiated samples were dominated by Firmicutes and Bacteroidetes with Firmicutes being significantly more abundant than Bacteroidetes. In the mucosal compartment of the Vehicle UI group, Firmicutes contributed 73.62 ± 0.07% and Bacteroidetes 24.30 ± 0.06% and in the MIIST305 UI group, Firmicutes represented 70.24 ± 0.07%, while Bacteroidetes accounted for 26.90 ± 0.07%. The abundance of Firmicutes at 6 DPI decreased in both Vehicle IR and MIIST305 IR groups, although this reduction was not statistically significant. Other phyla such as Verrucomicrobia, Actinobacteria, and Tenericutes were also present in the luminal and mucosal samples albeit at much lower relative abundance. Radiation exposure led to a dramatic increase in the relative abundance of *Verrucomicrobia*. In particular, samples from Vehicle IR exhibited an average relative abundance of 38.24 ± 0.03 % in the luminal compartment and 22.72 ± 0.08 % in the mucosal compartment at 6 DPI. In contrast, in Vehicle UI samples, the relative abundance of *Verrucomicrobia* was markedly lower, with values of 0.48 ± 0.01% in the luminal and 0.22 ± 0.01 % in the mucosal compartments. A similar increase was noted between the unirradiated and 6 DPI MIIST305-treated groups, with a 40.56 ± 0.07 % rise in luminal samples and a 28.10 ± 0.08% increase in mucosal samples. On the other hand, there was a significant reduction in relative abundance of Bacteroidetes and Actinobacteria, whereas Tenericutes were more abundant in the mucosal compartment under basal condition and displayed a differential response to radiation with a significant reduction in their abundance in the luminal compartment. In contrast, there was a marked enrichment of Tenericutes (Figure S4a) in the mucosal compartment, especially in Vehicle treated group by 63.35 ± 52.74 % (p = 2.0E-03) with respect to MIIST305 IR group. Taxonomic analysis at the genus levels revealed that *Anaeroplasma* (Figure S4b) accounted entirely for the relative abundance of Tenericutes. MIIST305 treated bacterial communities displayed the same composition as the vehicle-treated communities. Differential abundance of the microbiome at the genus level revealed a reduction in beneficial bacteria such as *Alistipes*, *Eubacterium*, *Lactobacillus*, and *Bifidobacterium* (Figure 6k) and an increase of potentially pathogenic bacteria like *Enterococcus* and *Staphylococcus* (Figure 6l) in both the luminal and mucosal samples of irradiated animals compared with unirradiated animals, whereas MIIST305 treatment did not have an impact on these microbial shifts at 6 DPI.

Next, we compared the microbiome diversity and composition of Vehicle IR and MIIST305 IR samples on 12 DPI. α -diversity using the Chao1 index showed that MIIST305 IR genera richness was significantly higher in both luminal (p = 1.53E-02) and mucosal (p = 1.04E-02) samples compared with Vehicle IR samples (Figures 7a and 7b). In contrast, Shannon index did not show appreciable differences between the two groups (Figures 7c and 7d). β-diversity examination using the Bray-Curtis dissimilarity test and visualized by PCoA showed that MIIST305 IR samples clustered separately from Vehicle IR samples in both lumen and mucosa (Figures 7e and 7f), implying the two groups had significantly different bacterial composition. The relative abundance at the phylum rank showed that Firmicutes abundance had mostly recovered from 6 DPI and almost completely dominated the phylum taxa, whereas Bacteriodetes remained at low levels. There was a significant reduction in Verrucomicrobia abundance in both luminal and mucosal samples, especially in the MIIST305 IR samples (Figures 7g, 7h, 7i). Among differentially abundant bacteria at genus level, presence of *Alistipes, Acetitomaculum, Eubacterium* and *Bifidobacterium* were highly enriched in MIIST305 IR samples with respect to Vehicle IR. In contrast, the abundance of *Enterococcus* and *Staphylococcus* in luminal and mucosal samples from MIIST305 samples were significantly reduced compared with Vehicle IR. Finally, a composition analysis at the species levels revealed that *Lactobacillus johnsonii* abundance accounted for the majority of species (Figures 4c and 4d) and it was more abundant in the MIIST305 IR samples compared with Vehicle IR samples. The relative abundance of *L. johnsonii* was 33.7 ± 27.64 % vs 64.3 ± 6.83 % (p = 4.29E-02) in the luminal Vehicle IR vs MIIST305 IR vs and 16.7 ± 13.19 % vs 42.3 ± 16.01 % (p = 2.50E-02) in the mucosal compartment of MIIST305 IR vs. Vehicle IR.

**Figure 7.**
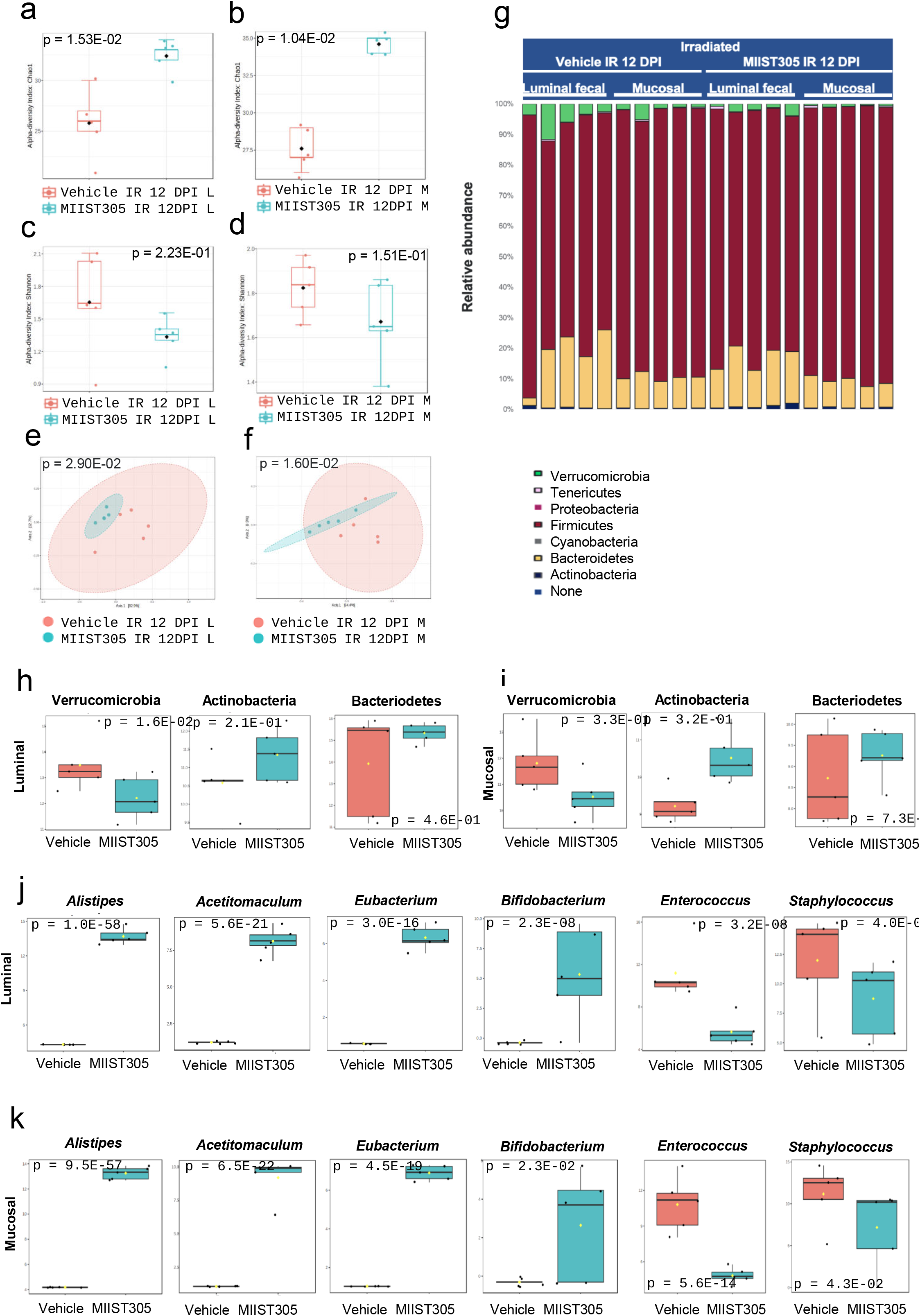
MIIST305 reduces x -ray induced gut dysbiosis in C57BL/6 (n=5) mice at 12 DPI. Representation of Chao1index (a) luminal, (b) mucosal, Shanon index (c) luminal, (d) mucosal of the intestinal microbiota between Vehicle IR 12DPI and MIIST305 IR 12DPI. PCoA score plots (Bray– Curtis) between Vehicle and MIIST305 at genus level (e) Luminal (f) Mucosal. Statistical analysis was performed using the PERMANOVA test, with p < 5.0E-02 considered statistically significant, (g) relative abundance at phylum level, differential abundance of bacteria at phylum level (h) luminal (i) mucosal, differential abundance of bacterial genera (j) luminal (k) mucosal, L and M denotes luminal and mucosal, respectively. For α-diversity (Chao1 and Shannon index) and β-diversity (PCoA plots) analysis, comparison between test groups was conducted using the Mann-Whitney U test and the PERMANOVA test respectively. For relative abundance at phylum level and differential abundance at genus levels, comparisons between treatment groups were performed using two-tailed unpaired t tests, the data are presented as mean ± SEM and p < 5.0E- 02 is considered statistically significant.

## Discussion

As a rapidly self-renewing tissue, the GI epithelium is particularly sensitive to ionizing radiation (IR)^54–57^. Radiation-induced intestinal epithelial cell death results in the loss of GI epithelial integrity and function characterized by epithelial erosion, crypt distortion^58^, loss of granularity^59^ and goblet cell depletion^60^. These changes compromise the gut barrier allowing luminal antigens and microorganisms to penetrate the lamina propria and enter the bloodstream through the MLN. This breach triggers a strong local and systemic pro-inflammatory response, which further exacerbates damage to the gut^61^.

Our results show that MIIST305 exhibits strong radio-mitigating effects in mice, with treatment administered 24 hours, after a single high dose of partial body irradiation followed by additional doses on days 3 and 5 resulted in approximately 85% survival by day 30, whereas all vehicle-treated animals died within 10 days due to GI injury^62–63^. We show that radiation exposure reduced the number of proliferating^64^ cells and epithelial goblet cells^65^ in the colon compared with unirradiated mice. However, we found that treatment with MIIST305 significantly prevented shortening of colon length at 12 DPI compared to vehicle-treated animals. Further, an increased number of proliferating cells per colonic crypt was also observed in comparison to vehicle-treated animals, accompanied with a markedly increased number of goblet cells per crypt as well as a higher production of mucins per cells, on both days 6 and 12 post-irradiation, as measured by Alcian Blue staining suggesting that MIIST305 could play an important role in maintaining the integrity of the mucus layer after radiation exposure and reinforces the gut barrier function. Concordantly, we show that MIIST305 treatment reduced gut barrier permeability and greatly diminished bacterial translocation to MLN.

In the present work, we also measured the concentration of a panel of cytokines in serum and colon lysates that have been associated with GI injury. The data show that pro-inflammatory cytokines were reduced by MIIST305 treatment in the serum compared with vehicle-treated irradiated animals. Most of the proinflammatory cytokines that were upregulated by radiation and reduced by MIIST305 were related to macrophage function with roles in trafficking immune cells^66–69^. In contrast, the anti-inflammatory cytokine IL-10^70^ was significantly upregulated in the serum of the MIIST305-treated mice compared with vehicle-treated mice. We observed that most of the serum cytokines that were upregulated in the serum, were also upregulated in the colon, although MIIST305 had a limited impact on colonic cytokines. We demonstrate that MIIST305 transiently reduced levels of IL-6 and increased levels of IL-33. IL-6 exerts a dual role on GI physiology, which is modulated by the composition of gut microbiota. During bacterial dysbiosis, IL-6 has been demonstrated to promote inflammatory responses and tissue damage^71^. IL-33 is an alarmin cytokine that is released upon tissue damage^72^ and it has been shown to restore barrier integrity by promoting IEC regeneration after intestinal injury and increase goblet cell numbers and Muc2 production^73^. Furthermore, IL-33 promotes an immunosuppressive environment in the presence of IL-10 by inducing regulatory T cell function in the intestine^74^. Recently, it has been shown that IL-33 potentiates intestinal stem cell regeneration in ileum after radiation injury through a mechanism that involves secretion of EGF by Paneth cells^75^. We now show that radiation exposure results in a robust induction of IL-33 in the mouse colon as well as that MIIST305 upregulates its expression. Considering that Paneth cells are not normally found in colon, it would be of interest to elucidate the mechanism of IL-33 induction in the colon in response to irradiation.

We explored the impact of radiation on the gut microbiome community in both the luminal and colonic mucosal compartment in mice treated with MIIST305 or vehicle by measuring α- and β-diversity. In the human gut, it has been demonstrated that the luminal microbiota and the mucosa-associated microbiota differ in diversity and composition^76–77^. Radiation exposure reduced α-diversity (Chao1 index) on day 6 post-irradiation, whereas irradiated and unirradiated samples were clustered separately (β-diversity) suggesting the two bacterial communities were significantly different from each other. MIIST305 treatment did not have a significant effect on α- diversity at day 6. By day 12, the MIIST305-treated mice indicated greater α-diversity compared with vehicle-treated animals. In contrast, β-diversity analysis demonstrated a clear separation between unirradiated and irradiated microbiota groups in both the lumen and the mucosa at day 6 in the Vehicle and MIIST305-treated mice on y day 12, implying the two groups had divergent microbial composition.

Taxonomic profiling of gut microbiota at the phylum level revealed that the abundance of Firmicutes did not change appreciably before and after irradiation; however, Bacteroidetes abundance was greatly diminished at both days 6 and 12 following irradiation. Although radiation caused a large increase in the relative abundance of Verrucomicrobia (represented by the species *Akkermansia muciniphila*) at day 6 which is consistent with previous reports^78–80^, we show here that this increase was transient and greatly diminished by day 12 post-irradiation, especially in the MIIST305-treated animals that returned to pre-irradiation levels. *A. muciniphila* is a probiotic under homeostatic conditions and has been associated with several beneficial effects^81^ but it has also been associated with adverse effects in some circumstances. For example, increased levels of *A. muciniphila* have been reported in colitis patients and it is negatively associated with survival after total body irradiation^82^.

Besides Verrucomicrobia, the relative abundance of Bacteroidetes and Actinobacteria was reduced following irradiation in both the lumen and colonic mucosa similar to previous findings^83^. On the other hand, the abundance of Tenericutes in the colonic mucosa increased following radiation exposure. Previous studies have reported that elevated levels of Tenericutes are associated with chronic gut inflammation and are directly linked to increased expression of IL-6 in the colonic environment^84^, a pattern consistent with the findings of our study. MIIST305-treated mice exhibited a reduced abundance of Tenericutes in the mucosal compartment, along with a significant decrease in IL-6 levels at 6DPI, indicating a suppression of inflammation.

A number of beneficial bacterial genera were also driving the differences on day 6 after irradiation. Specifically, *Alistipes*, *Bifidobacterium*, *Eubacterium*, and *Lactobacillus* levels were severely reduced compared with unirradiated mice. It is thought that the genus *Alistipes* may have a protective role against colitis and has been found to be suppressed in response to radiation and chemotherapy and has been suggested to mitigate radiation injury^77.^ *Bifidobacterium spp*. are known probiotics that increase production of Muc2 and tight junction proteins, thus, decreasing intestinal permeability and reducing bacterial translocation. In relation to human disease, *Bifidobacterium* spp. have been shown to downregulate Toll-like receptor 4 (TLR4)/NF-κB signaling in IECs and prevent acute colitis. *Eubacterium* spp. are major butyrate producers in the gut that have beneficial impact on colonocytes and the intestinal epithelial barrier. They suppress TNFα-induced TLR4/NF-κB axis and intestinal inflammation^85^. They also produce valerate that has been shown to confer radioresistance^13^. The genera *Enterococcus* and *Staphylococcus*, which include several potentially pathogenic species, were nearly undetectable in animals under homeostatic conditions. However, radiation exposure led to a marked increase in the abundance of these genera in both vehicle- and MIIST305-treated mice at 6DPI. MIIST305 treatment significantly modulated the microbiome, inducing notable shifts in several genera. Specifically, MIIST305-treated mice exhibited a pronounced increase in *Alistipes*, *Acetatimaculum*, *Eubacterium*, and *Bifidobacterium* compared to vehicle-treated mice. In contrast, *Enterococcus* and *Staphylococcus* levels were significantly reduced in MIIST305-treated mice relative to vehicle-treated animals, though they remained elevated compared to unirradiated controls.

Finally, *Lactobacillus* was suppressed on day 6 after irradiation. However, *Lactobacillus* specifically *Lactobacillus johnsonii* was the most prevalent *Lactobacillus* species driving alterations on day 12 and its levels were significantly higher in the MIIST305-treated gut microbiota. *Lactobacillus johnsonii* ameliorates oxidative stress, systemic genotoxicity^78^, and alleviates the damage caused by microbial pathogens by inhibiting TLR4, NF-κB, and the inflammasome (NLRP3) signaling pathway^86^. *L. johnsonii* is an acetate producer ^87^ and promotes gut barrier homeostasis. Furthermore, combination of *Lactobacillus* and *Bifidobacterium* are implicated in the synthesis of secondary bile acids that show robust anti-inflammatory properties^88^.

In summary, we show that MIIST305 has significant potential as an effective mitigator of GI-ARS. MIIST305 enhances the survival rate of irradiated mice by preserving the intestinal barrier function, reducing bacterial translocation, suppressing radiation-induced inflammatory cytokine storm, ultimately facilitating the recovery of the structure and function of the intestine. Furthermore, MIIST305 reduces systemic inflammation in response to radiation suggesting that the drug could alleviate the risk of detrimental immune-mediated organ damage. Whereas radiation exposure leads to bacterial microbial imbalance (dysbiosis), which is characterized by the loss of beneficial species and the overgrowth of pathobionts, MIIST305 decreases potential pathobionts while expanding commensal protective bacteria. Future studies are directed towards determining the mechanism of action of MIIST305, as well as providing the mechanistic relationship between microbiota and radiation-induced bowel injury. Associative studies between gut microbiota composition, histopathological alterations, and cytokine modulation may enlighten specific microbial taxa with a causative impact on GI-ARS. Furthermore, elucidating significantly changed circulating and tissue cytokines after partial body irradiation will aid in deciphering the mechanism of GI-ARS and identifying potential biomarkers to predict clinical outcome.

## Abbreviations

BM5, 5% bone marrow shielding; CFU, Colony forming units; CUIMC, Columbia University Irving Medical Center; FMT, fecal microbiota transplant; GI, gastrointestinal; IECs, intestinal epithelial cells; MCMs, medical countermeasures; MIIST, multivalent innate immune signaling target platform; PBI, partial body irradiation; SSD, source to surface distance; IACUC, Institutional Animal Care and Use Committee; UI, unirradiated; IR, irradiated; MISS, Mouse Interventional Scoring System; MLNs, mesenteric lymph nodes; PBS, phosphate-buffered saline ; PCoA, Principal coordinates analysis ; PERMANOVA, Permutational multivariate analysis of variance; WBC, white blood cells.

## Acknowledgements

The authors would like to thank the Molecular Pathology core facility for technical assistance with histology and immunohistochemistry

## Author contributions

**Debmalya Mitra:** Methodology, Investigation, Writing-Original Draft, Formal Analysis.: **Gabriel K. Armijo:** Methodology, Investigation, Writing-Review and Editing**.: Elizabeth H. Ober:** Investigation, Formal Analysis, Writing-Review and Editing.: **Shenda M. Baker**: Writing-Review and Editing, **Helen C. Turner:** Conceptualization, Funding acquisition, Writing- Review and Editing, Resources. **Constantinos G. Broustas:** Conceptualization, Methodology, Writing-Review and Editing, Supervision, Resources, Project administration.

## ORCID

**Debmalya Mitra:** https://orcid.org/0000-0003-1560-5662

**Gabriel K. Armijo:** https://orcid.org/0000-0002-7930-8248

**Elizabeth H. Ober:** https://orcid.org/0009-0004-5603-9462

**Shenda M. Baker**: https://orcid.org/0000-0002-9985-473X

**Helen C. Turner**: https://orcid.org/0000-0002-3087-4740

**Constantinos G. Broustas:** https://orcid.org/0000-0001-7169-807X

## Availability of Data

All the generated data from the study are either embedded in the main text file or in the supplementary files, any further details will be made available on request.

---

**Supplementary Figure 1.** Administration of MIIST305 enhances survival and prevents body weight loss in C57BL/6 mice exposed to 12.5 Gy PBI/BM5. (a) The Kaplan-Meier curve illustrates the survival of the treatment groups up to 12 DPI between treatment groups, Vehicle IR (n=20) and MIIST305 IR (n=8). Statistical analysis was performed by Log-rank (Mantel-Cox) test (b) Comparison of body weight loss (%) following exposure of X-ray at 12.5 Gy, PBI/BM5 measured till 12 DPI. Data are presented as mean ± SEM, two-tailed unpaired t-tests were conducted between treatment groups and *p < 5.0E-02 is considered statistically significant with respect to the Vehicle IR.

**Supplementary Figure 2.** Changes in the serum cytokine levels (pg/ml) in response to 12.5 Gy X ray irradiation in C57/BL6 mice from different treatment groups at 6 DPI and 12 DPI. (a) IL-1β, (b) IL-2, (c) IFN-γ, (d) IL-17A, (e) MIP-1 α, (f) IL-22, (g) IL-23, (h) IL-27p28/IL-30. Comparisons between treatment groups (n = 3 to 8/ group) were performed using two-tailed unpaired t-tests, and the data are presented as mean ± SEM.

**Supplementary Figure 3.** Changes in the cytokine levels in the colonic tissues of C57/BL6 mice from different treatment groups at 6 DPI and 12 DPI in response to 12.5Gy x -ray irradiation. (a) IL-1β, (b) IL-2, (c) TNF- α, (d) KC/GRO, (e ) IFN-γ, (f) IL-10, (g) IL-17A, (h) IL-17A/F, (i) IL-22, (j) IL-23, (k) IL-27p28/IL-30, (l) IP-10, (m) MIP-1α, (n) MIP-1β, (o) MIP-2, (p) MIP-3α, (q) MCP-1 Comparisons between treatment groups (n = 3 to 8/ group) were performed using two-tailed unpaired t-tests, and the data are presented as mean ± SEM.

**Supplementary Figure 4.** MIIST305 treatment reduces relative abundance of Tenericutes in the mucosal compartment of C57BL/6 mice at 6 DPI and increases abundance of *Lactobacillus johnsonii* at 12 DPI. % relative abundance of (a) Tenericutes, (b) *Anaeroplasma* (c) *L. johnsonii* at 6 DPI and (d) *L. johnsonii* at 12 DPI. L and M denotes luminal and mucosal, respectively. Comparisons between treatment groups were performed using two-tailed unpaired t-tests (n=5), and the data are presented as mean ± SEM. *P < 5.0E-02 is considered statistically significant.

## References

1. Winters TA, Marzella L, Molinar-Inglis O, et al. Gastrointestinal Acute Radiation Syndrome: mechanisms, models, markers, and medical countermeasures. Radiat Res. 2024;201(6):628–646. doi:10.1667/RADE-23-00196.1

2. MacVittie TJ, Farese AM. Defining the concomitant multiple organ injury within the ARS and DEARE in an animal model research platform. Health Phys. 2020;119(5):519–526. doi:10.1097/HP.0000000000001327

3. Hollingsworth BA, Cassatt DR, DiCarlo AL, et al. Acute radiation syndrome and the microbiome: impact and review. Front Pharmacol. 2021;12:643283. doi:10.3389/fphar.2021.643283

4. Singh VK, Seed TM. Radiation countermeasures for hematopoietic acute radiation syndrome: growth factors, cytokines and beyond. Int J Radiat Biol. 2021;97(11):1526–1547. doi:10.1080/09553002.2021.1969054

5. Hou K, Wu ZX, Chen XY, et al. Microbiota in health and diseases. Signal Transduct Target Ther. 2022;7(1):135. doi:10.1038/s41392-022-00974-4

6. Barbara G, Barbaro MR, Fuschi D, et al. Inflammatory and microbiota-related regulation of the intestinal epithelial barrier. Front Nutr. 2021;8:718356. doi:10.3389/fnut.2021.718356. [published correction appears in Front Nutr. 2021:790387. doi: 10.3389/fnut.2021.790387]

7. Maciel-Fiuza MF, Muller GC, Campos DMS, et al. Role of gut microbiota in infectious and inflammatory diseases. Front Microbiol. 2023;14:1098386. doi:10.3389/fmicb.2023.1098386

8. Cui M, Xiao H, Li Y, et al. Faecal microbiota transplantation protects against radiation-induced toxicity. EMBO Mol Med. 2017;9(4):448–461. doi:10.15252/emmm.201606932

9. Li M, Gu MM, et al. The vanillin derivative VND3207 protects intestine against radiation injury by modulating p53/NOXA signaling pathway and restoring the balance of gut microbiota. Free Radic Biol Med. 2019;145:223–236. doi: 10.1016/j.freeradbiomed.2019.09.035. Erratum in: Free Radic Biol Med. 2023;204:83. doi: 10.1016/j.freeradbiomed.2023.04.016. PMID: 31580946.

10. Kim YS, Kim J, Park SJ. High-throughput 16S rRNA gene sequencing reveals alterations of mouse intestinal microbiota after radiotherapy. Anaerobe. 2015;33:1–7. doi:10.1016/j.anaerobe.2015.01.004

11. Zhao Z, Cheng W, Qu W, et al. Antibiotic alleviates radiation-induced intestinal injury by remodeling microbiota, reducing inflammation, and inhibiting fibrosis. ACS Omega. 2020;5(6):2967–2977. doi:10.1021/acsomega.9b03906

12. Gerassy-Vainberg S, Blatt A, Danin-Poleg Y, et al. Radiation induces proinflammatory dysbiosis: transmission of inflammatory susceptibility by host cytokine induction. Gut. 2018;67(1):97–107. doi:10.1136/gutjnl-2017-313789

13. Li Y, Yan H, Zhang Y, et al. Alterations of the gut microbiome composition and lipid metabolic profile in radiation enteritis. Front Cell Infect Microbiol. 2020;10:541178. doi:10.3389/fcimb.2020.541178

14. Goudarzi M, Mak TD, Jacobs JP, et al. An integrated multi-omic approach to assess radiation injury on the host-microbiome axis. Radiat Res. 2016;186(3):219–234. doi:10.1667/RR14306.1

15. Zhao Y, Zhang J, Han X, Fan S. Total body irradiation induced mouse small intestine senescence as a late effect. J Radiat Res. 2019;60(4):442–450. doi:10.1093/jrr/rrz026

16. Guo H, Chou WC, Lai Y, et al. Multi-omics analyses of radiation survivors identify radioprotective microbes and metabolites. Science. 2020; 370(6516):eaay9097.

17. Fernandes A, Oliveira A, Soares R, et al. The Effects of ionizing radiation on gut microbiota: what can animal models tell us?-a systematic review. Curr Issues Mol Biol. 2023;45(5):3877–3910. doi:10.3390/cimb45050249

18. Lu L, LiF, Gao Y. et al. Microbiome in radiotherapy: an emerging approach to enhance treatment efficacy and reduce tissue injury. Mol Med. 2024 30, 105. 10.1186/s10020-024-00873-0

19. Gieryńska M, Szulc-Dąbrowska L, Struzik J, et al. Integrity of the intestinal barrier: the involvement of epithelial cells and microbiota-a mutual relationship. Animals (Basel). 2022;12(2):145. doi:10.3390/ani12020145

20. Capaldo CT, Powell DN, Kalman D. Layered defense: how mucus and tight junctions seal the intestinal barrier. J Mol Med (Berl). 2017;95(9):927–934. doi:10.1007/s00109-017-1557-x

21. Kong C, Elderman M, Cheng L, et al. Modulation of intestinal epithelial glycocalyx development by human milk oligosaccharides and non-digestible carbohydrates. Mol Nutr Food Res. 2019;63(17):e1900303. doi:10.1002/mnfr.201900303

22. Capaldo CT, Nusrat A. Claudin switching: Physiological plasticity of the Tight Junction. Semin Cell Dev Biol. 2015;42:22–29. doi:10.1016/j.semcdb.2015.04.003

23. Cornick S, Tawiah A, Chadee K. Roles and regulation of the mucus barrier in the gut. Tissue Barriers. 2015;3(1-2):e982426. doi:10.4161/21688370.2014.982426.

24. Sun J, Shen X, Li Y, et al. Therapeutic potential to modify the mucus barrier in inflammatory bowel disease. Nutrients. 2016;8(1):44. doi:10.3390/nu8010044

25. Johansson ME, Larsson JM, Hansson GC. The two mucus layers of colon are organized by the MUC2 mucin, whereas the outer layer is a legislator of host-microbial interactions. Proc Natl Acad Sci U S A. 2011;108 Suppl 1(Suppl 1):4659-4665. doi:10.1073/pnas.1006451107

26. Ma Y, Huang Z, Li C. The bile acids specifically modulate colonic MUC2 and tight junction protein expression in the human colon cancer cell line. Curr Dev Nutr. 2021;5(Suppl 2):33. doi:10.1093/cdn/nzab033_033

27. Zhang M, Wu C. The relationship between intestinal goblet cells and the immune response. Biosci Rep. 2020;40(10):BSR20201471. doi:10.1042/BSR20201471

28. Derrien M, van Passel MW, van de Bovenkamp JH, et al. Mucin-bacterial interactions in the human oral cavity and digestive tract. Gut Microbes. 2010;1(4):254–268. doi:10.4161/gmic.1.4.12778

29. Van der Sluis M, De Koning BA, De Bruijn AC, et al. Muc2-deficient mice spontaneously develop colitis, indicating that MUC2 is critical for colonic protection. Gastroenterology. 2006;131(1):117–129. doi:10.1053/j.gastro.2006.04.020

30. Sterling JD and Baker SM. Electro-lyotropic equilibrium and the utility of ion-pair dissociation constants. Colloid and Interface Science Comm 2017;20, 9-11.

31. Sterling JD and Baker SM. A continuum model of mucosa with glycan-ion pairing. Macromolecular Theory and Simulation 2018;doi/10.1002/mats.201700079.

32. Cui W, Hull L, Zizzo A, et al. The Roles of IL-18 in a realistic partial body irradiation with 5% bone marrow sparing (PBI/BM5) model. Toxics 2024, 12, 5. 10.3390/toxics12010005

33. Kumar VP, Wuddie K, Tsioplaya A, et al. Development of a multi-organ radiation injury model with precise dosimetry with focus on GI-ARS. Radiat Res. 2024;201(1):19–34. doi:10.1667/RADE-23-00068.1

34. Fish BL, MacVittie TJ, Gao F, et al. Rat models of partial-body irradiation with bone marrow-sparing (leg-out PBI) designed for FDA approval of countermeasures for mitigation of acute and delayed injuries by radiation. Health Phys. 2021;121(4):419–433. doi:10.1097/HP.0000000000001444

35. Booth C, Tudor G, Tudor J, et al. Acute gastrointestinal syndrome in high-dose irradiated mice. Health Phys. 2012;103(4):383–399. doi:10.1097/hp.0b013e318266ee13

36. Plett PA, Sampson CH, Chua HL, et al. Establishing a murine model of the hematopoietic syndrome of the acute radiation syndrome. Health Phys. 2012;103(4):343–355. doi:10.1097/HP.0b013e3182667309

37. Koch A, Gulani J, King G, et al. Establishment of early endpoints in mouse total-body irradiation model. PLoS One. 2016;11(8):e0161079. Published 2016 Aug 31. doi:10.1371/journal.pone.0161079

38. Rose WA 2nd, Sakamoto K, Leifer CA. TLR9 is important for protection against intestinal damage and for intestinal repair. Sci Rep. 2012;2:574. doi:10.1038/srep00574

39. Breynaert C, Dresselaers T, Perrier C, et al. Unique gene expression and MR T2 relaxometry patterns define chronic murine dextran sodium sulphate colitis as a model for connective tissue changes in human Crohn’s disease. PLoS One. 2013;8(7):e68876. doi:10.1371/journal.pone.0068876

40. Grillo F, Campora M, Pigozzi S, et al. Methods for restoration of ki67 antigenicity in aged paraffin tissue blocks. Histochem Cell Biol. 2021;156(2):183–190. doi:10.1007/s00418-021-01987-w

41. Lei HT, Yan S, He YH, et al. Ki67 testing in the clinical management of patients with non-metastatic colorectal cancer: Detecting the optimal cut-off value based on the Restricted Cubic Spline model. Oncol Lett. 2022;24(6):420. doi:10.3892/ol.2022.13540

42. Bagalagel A, Diri R, Noor A, et al. Curative effects of fucoidan on acetic acid induced ulcerative colitis in rats via modulating aryl hydrocarbon receptor and phosphodiesterase-4. BMC Complement Med Ther. 2022;22(1):196. doi:10.1186/s12906-022-03680-4

43. Peng YJ, Shen TL, Chen YS, et al. Adiponectin and adiponectin receptor 1 overexpression enhance inflammatory bowel disease. J Biomed Sci. 2018;25(1):24. doi:10.1186/s12929-018-0419-3

44. Yang, J., Elbaz-Younes, I., Primo, C. et al. Intestinal permeability, digestive stability and oral bioavailability of dietary small RNAs. Sci Rep 8, 10253 (2018). doi: 10.1186/s12263-017-0563-5

45. Kang M, Mischel RA, Bhave S, et al. The effect of gut microbiome on tolerance to morphine mediated antinociception in mice. Sci Rep. 2017; 7:42658. doi:10.1038/srep42658

46. Choi Y, Lichterman JN, Coughlin LA, et al. Immune checkpoint blockade induces gut microbiota translocation that augments extraintestinal antitumor immunity. Sci Immunol. 2023;8(81):eabo2003. doi:10.1126/sciimmunol.abo2003

47. Buchta Rosean C, Bostic RR, Ferey JCM, et al. Preexisting commensal dysbiosis is a host-intrinsic regulator of tissue inflammation and tumor cell dissemination in hormone receptor-positive breast cancer. Cancer Res. 2019;79(14):3662–3675. doi:10.1158/0008-5472.CAN-18-3464

48. Callahan BJ, McMurdie PJ, Rosen MJ, et al. DADA2: High-resolution sample inference from Illumina amplicon data. Nat Methods. 2016;13(7):581–583. doi:10.1038/nmeth.3869

49. Lu Y, Zhou G, Ewald J, et al. MicrobiomeAnalyst 2.0: comprehensive statistical, functional and integrative analysis of microbiome data. Nucleic Acids Res. 2023;51(W1): W310–W318. doi:10.1093/nar/gkad407

50. Takada Y, Hisamatsu T, Kamada N, et al. Monocyte Chemoattractant Protein-1 contributes to gut homeostasis and intestinal inflammation by composition of IL-10– producing regulatory macrophage subset. J Immunol 1 2010;184 (5): 2671–2676. 10.4049/jimmunol.0804012.

51. Shukla PK, Meena AS, Gangwar R, et al. LPAR2 receptor activation attenuates radiation-induced disruption of apical junctional complexes and mucosal barrier dysfunction in mouse colon. FASEB J. 2020;34(9):11641–11657. doi:10.1096/fj.202000544R

52. Hiippala K, Jouhten H, Ronkainen A, et al. The potential of gut commensals in reinforcing intestinal barrier function and alleviating inflammation. Nutrients. 2018;10(8):988. doi:10.3390/nu10080988

53. Yu Y, Lin X, Feng F, et al. Gut microbiota and ionizing radiation-induced damage: Is there a link?. Environ Res. 2023;229:115947. doi:10.1016/j.envres.2023.115947

54. Novak JM, Collins JT, Donowitz M, et al. Effects of radiation on the human gastrointestinal tract. J Clin Gastroenterol. 1979;1(1):9–39. doi:10.1097/00004836-197903000-00003

55. Somosy Z, Horváth G, Telbisz A, et al. Morphological aspects of ionizing radiation response of small intestine. Micron. 2002;33(2):167–178. doi:10.1016/s0968-4328(01)00013-0

56. Clevers H. The intestinal crypt, a prototype stem cell compartment. Cell. 2013;154(2):274–284. doi:10.1016/j.cell.2013.07.004

57. Williams JM, Duckworth CA, Burkitt MD, et al. Epithelial cell shedding and barrier function: a matter of life and death at the small intestinal villus tip. Vet Pathol. 2015;52(3):445–455. doi:10.1177/0300985814559404

58. Mitra D, Sikdar S, Chakraborty M, et al. Gum Odina prebiotic prevents experimental colitis in C57BL/6 mice model and its role in shaping gut microbial diversity, Food Bioscience. 53, 2023, 102509,

59. Cader MZ, Kaser A. Recent advances in inflammatory bowel disease: mucosal immune cells in intestinal inflammation. Gut. 2013;62(11):1653–1664. doi:10.1136/gutjnl-2012-303955

60. Birchenough GM, Johansson ME, Gustafsson JK, et al. New developments in goblet cell mucus secretion and function. Mucosal Immunol. 2015;8(4):712–719. doi:10.1038/mi.2015.32

61. Bhanja P, Norris A, Gupta-Saraf P, et al. BCN057 induces intestinal stem cell repair and mitigates radiation-induced intestinal injury. Stem Cell Res Ther. 2018;9(1):26. Published 2018 Feb 2. doi:10.1186/s13287-017-0763-3

62. Kenchegowda D, Bolduc DL, Kurada L, et al. Severity scoring systems for radiation-induced GI injury - prioritization for use of GI-ARS medical countermeasures. Int J Radiat Biol. 2023;99(7):1037–1045. doi:10.1080/09553002.2023.2210669

63. Rotolo J, Stancevic B, Zhang J, et al. Anti-ceramide antibody prevents the radiation gastrointestinal syndrome in mice. J Clin Invest. 2012;122(5):1786–1790. doi:10.1172/JCI59920

64. Orzechowska-Licari EJ, LaComb JF, Giarrizzo M, et al. Intestinal epithelial regeneration in response to ionizing irradiation. J Vis Exp. 2022;(185):10.3791/64028. Published 2022 Jul 27. doi:10.3791/64028

65. Jameus A, Dougherty J, Narendrula R, et al. Acute impacts of ionizing radiation exposure on the gastrointestinal tract and gut microbiome in mice. Int J Mol Sci. 2024;25(6):3339. doi:10.3390/ijms25063339.

66. Grimm MC, Doe WF. Chemokines in Inflammatory Bowel Disease Mucosa: Expression of RANTES, Macrophage Inflammatory Protein (MIP)-1α, MIP-1β, and γ-Interferon-Inducible Protein-10 by Macrophages, Lymphocytes, Endothelial Cells, and Granulomas. Inflamm Bowel Dis. 1996;2(2):88–96

67. Ohtsuka Y, Lee J, Stamm DS, Sanderson IR. MIP-2 secreted by epithelial cells increases neutrophil and lymphocyte recruitment in the mouse intestine. Gut. 2001;49(4):526–533. doi:10.1136/gut.49.4.526

68. Katchar K, Kelly CP, Keates S, et al. MIP-3alpha neutralizing monoclonal antibody protects against TNBS-induced colonic injury and inflammation in mice. Am J Physiol Gastrointest Liver Physiol. 2007;292(5):G1263–G1271. doi:10.1152/ajpgi.00409.2006

69. Deng Z, Wang S, Wu C, et al. IL-17 inhibitor-associated inflammatory bowel disease: A study based on literature and database analysis. Front Pharmacol. 2023;14:1124628. doi:10.3389/fphar.2023.1124628

70. Wei H-X, Wang B and Li B. IL-10 and IL-22 in Mucosal immunity: driving protection and pathology. Front. Immunol. 2020;11:1315. doi: 10.3389/fimmu.2020.01315.

71. Shahini A, Shahini A. Role of interleukin-6-mediated inflammation in the pathogenesis of inflammatory bowel disease: focus on the available therapeutic approaches and gut microbiome. J Cell Commun Signal. 2023;17(1):55–74. doi:10.1007/s12079-022-00695-x

72. Seo, D.H., Che, X., Kwak, M.S. et al. Interleukin-33 regulates intestinal inflammation by modulating macrophages in inflammatory bowel disease. Sci Rep 7, 851 (2017). 10.1038/s41598-017-00840-2.

73. Wang Y, He C, Xin S, et al. A Deep view of the biological property of interleukin-33 and its dysfunction in the gut. Int J Mol Sci. 2023;24(17):13504. doi:10.3390/ijms241713504.

74. Sattler S, Ling GS, Xu D, et al. IL-10-producing regulatory B cells induced by IL-33 (Breg(IL-33)) effectively attenuate mucosal inflammatory responses in the gut. J Autoimmun. 2014;50(100):107–122. doi:10.1016/j.jaut.2014.01.032

75. Calafiore M, Fu YY, Vinci P, et al. A tissue-intrinsic IL-33/EGF circuit promotes epithelial regeneration after intestinal injury. Nat Commun. 2023;14(1):5411. doi:10.1038/s41467-023-40993-5

76. Ringel Y, Maharshak N, Ringel-Kulka T, et al. High throughput sequencing reveals distinct microbial populations within the mucosal and luminal niches in healthy individuals. Gut Microbes. 2015;6(3):173–181. doi:10.1080/19490976.2015.1044711

77. Cui W, Hull L, Zizzo A, et al. The gut microbiome changes in wild type and IL-18 knockout mice after 9.0 Gy total body irradiation. Anim Microbiome. 2023;5(1):42. doi:10.1186/s42523-023-00262-8

78. Horseman TS, Parajuli B, Frank AM, et al. MICROBIOME AND INFLAMMASOME ALTERATIONS FOUND DURING RADIATION DOSE FINDING IN A SINCLAIR MINIPIG MODEL OF GASTROINTESTINAL ACUTE RADIATION SYNDROME. Shock. 2024;62(4):556–564. doi:10.1097/SHK.0000000000002422

79. Kumagai T, Rahman F, Smith AM. The Microbiome and radiation induced-bowel injury: evidence for potential mechanistic role in disease pathogenesis. Nutrients. 2018;10(10):1405. doi:10.3390/nu10101405

80. Li Y, Yan H, Zhang Y, et al. Alterations of the gut microbiome composition and lipid metabolic profile in radiation enteritis. Front Cell Infect Microbiol. 2020;10:541178. doi:10.3389/fcimb.2020.541178

81. He KY, Lei XY, Wu DH, et al. *Akkermansia muciniphila* protects the intestine from irradiation-induced injury by secretion of propionic acid. Gut Microbes. 2023;15(2):2293312. doi:10.1080/19490976.2023.2293312

82. Li K, Epperly MW, Barreto GA, et al. Longitudinal fecal microbiome study of total body irradiated mice treated with radiation mitigators identifies bacterial associations with survival. Front Cell Infect Microbiol. 2021;11:715396. doi:10.3389/fcimb.2021.715396

83. Kumagai T, Rahman F, Smith AM. The microbiome and radiation induced-bowel injury: Evidence for potential mechanistic role in disease pathogenesis. Nutrients. 2018;10(10):1405. doi:10.3390/nu10101405

84. Potrykus M, Czaja-Stolc S, Stankiewicz M, et al. Intestinal microbiota as a contributor to chronic inflammation and its potential modifications. Nutrients. 2021;13(11):3839. doi:10.3390/nu13113839

85. Abdulqadir R, Engers J, Al-Sadi R. Role of *Bifidobacterium* in modulating the intestinal epithelial tight junction barrier: current knowledge and perspectives. Curr Dev Nutr. 2023;7(12):102026. doi:10.1016/j.cdnut.2023.102026

86. Chen S, Li Y, Chu B, et al. *Lactobacillus johnsonii*L531 alleviates the damage caused by *Salmonella Typhimurium* via inhibiting TLR4, NF-κB, and NLRP3 inflammasome signaling pathways. Microorganisms. 2021;9(9):1983. doi:10.3390/microorganisms9091983

87. Wen, Y., Yang, L., Wang, Z. et al. Blocked conversion of *Lactobacillus johnsonii* derived acetate to butyrate mediates copper-induced epithelial barrier damage in a pig model. Microbiome 11, 218 (2023). 10.1186/s40168-023-01655-2.

88. Hamade DF, Espinal A, Yu J, et al. Lactobacillus reuteri releasing IL-22 (LR-IL-22) facilitates intestinal radioprotection for whole-abdomen irradiation (WAI) of ovarian cancer. Radiat Res. 2022;198(1):89–105. doi:10.1667/RADE-21-00224.1

