## Supplementary figures and images for "MIIST305 mitigates gastrointestinal acute radiation syndrome injury and ameliorates radiation-induced gut microbiome dysbiosis"

### Supplemental Figures

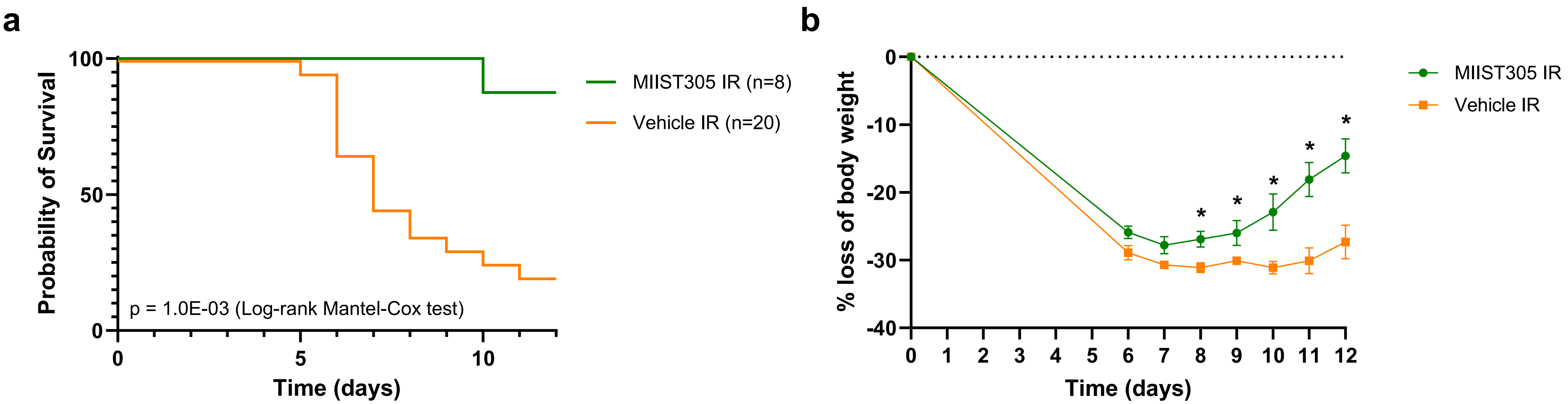

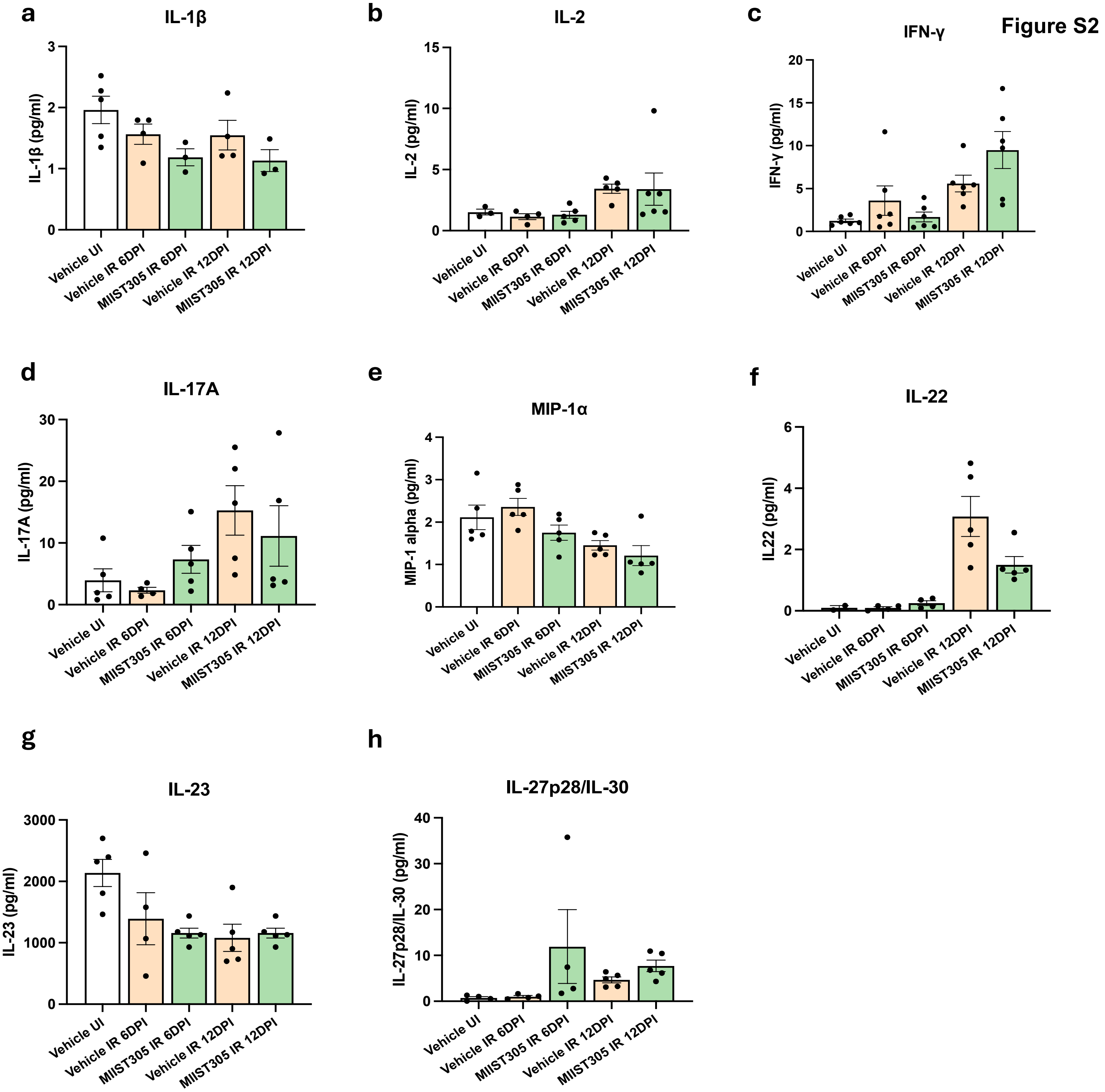

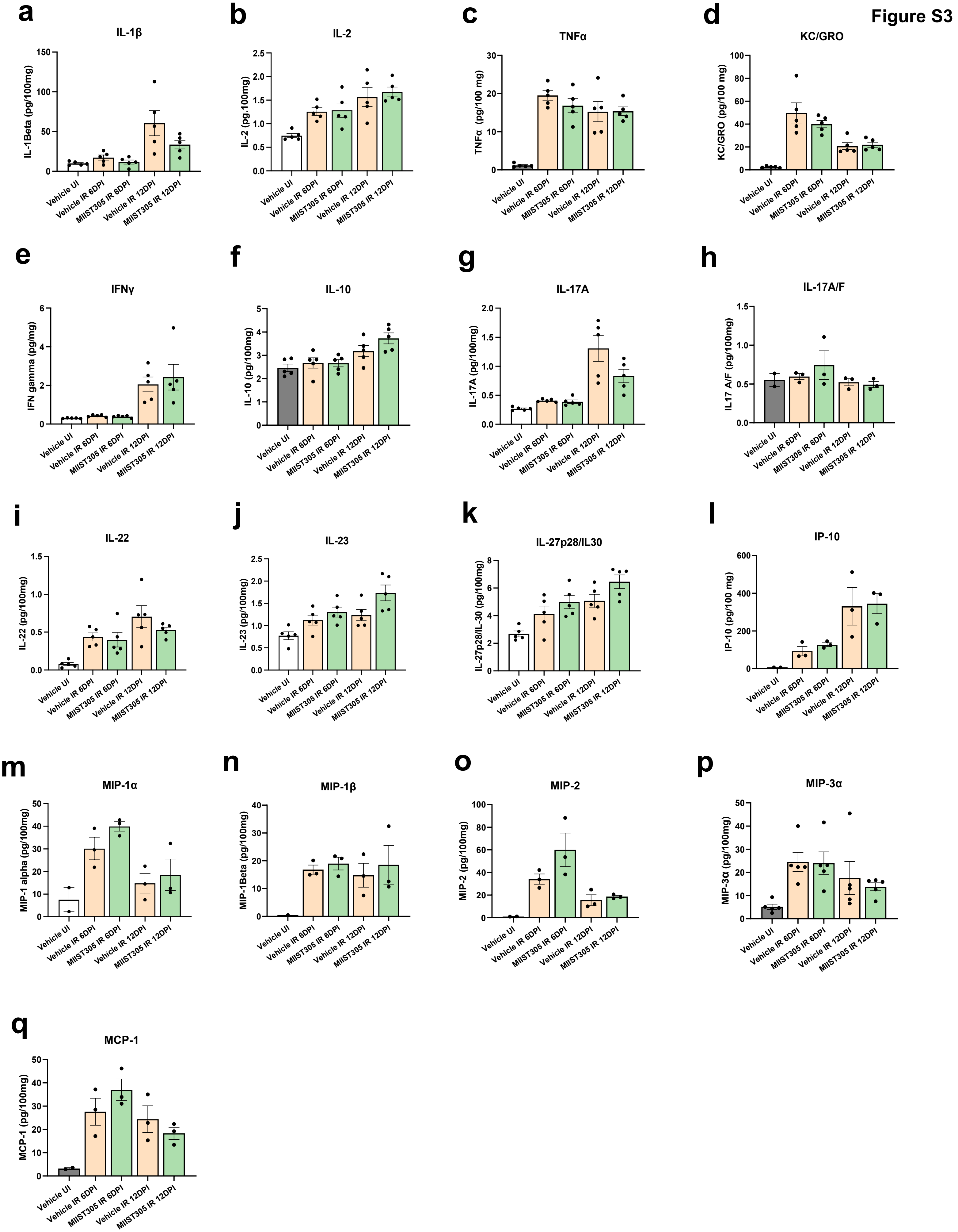

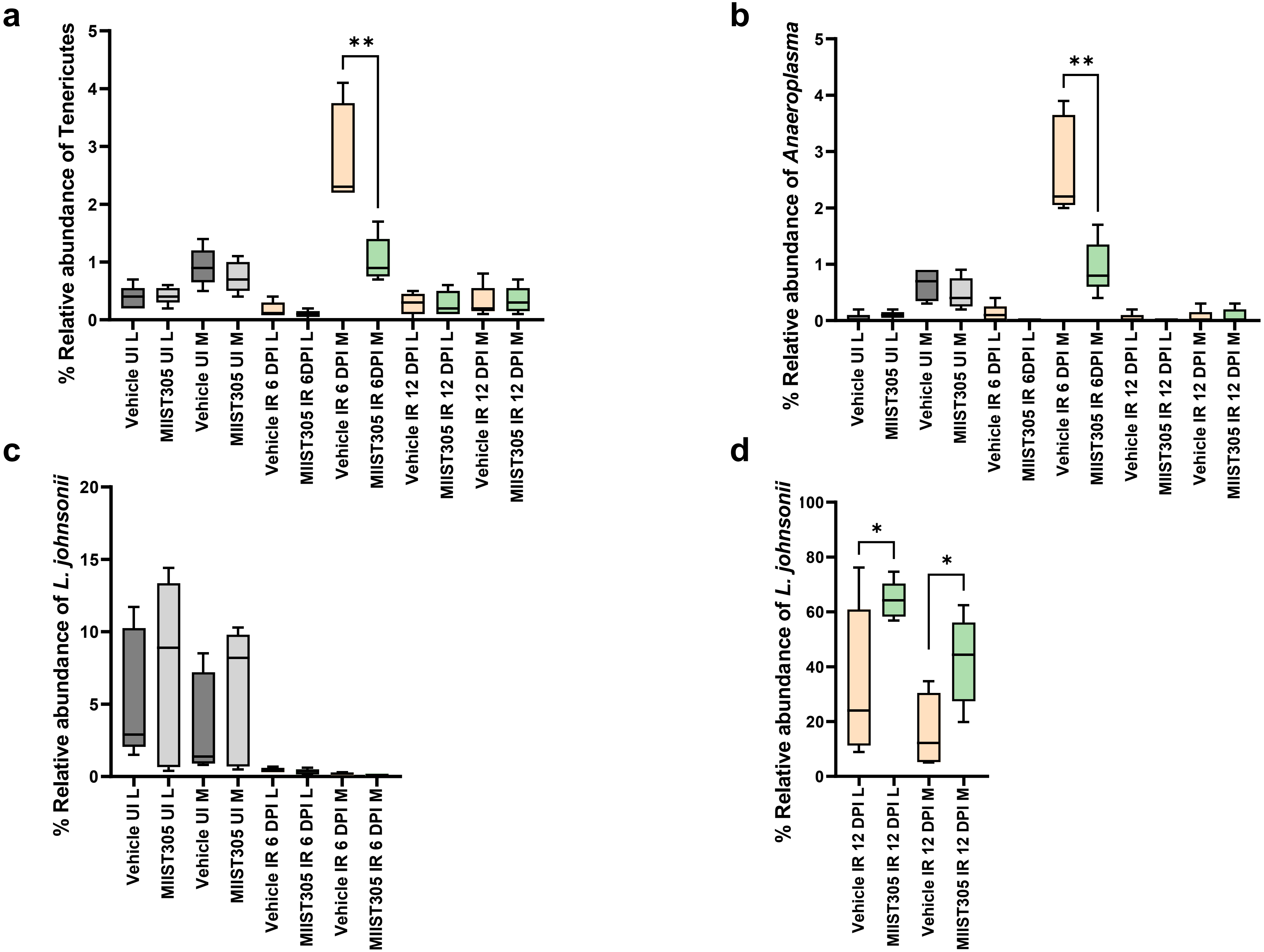
